# SMARCB1 loss interacts with neuronal differentiation state to block maturation and impact cell stability

**DOI:** 10.1101/2020.05.04.074443

**Authors:** Alison D. Parisian, Tomoyuki Koga, Shunichiro Miki, Pascal D. Johann, Marcel Kool, John R. Crawford, Frank B. Furnari

## Abstract

Atypical teratoid rhabdoid tumors (ATRT) are challenging pediatric brain cancers which are predominantly associated with inactivation of the gene SMARCB1, a conserved subunit of the chromatin remodeling BAF complex, which has known contributions to developmental processes. To identify potential interactions between SMARCB1 loss and the process of neural development, we introduced an inducible SMARCB1 loss of function system into human induced pluripotent stem cells (iPSCs) which were subjected to either directed neuronal differentiation or differentiation into cerebral organoids. Using this system, we have identified substantial differences in the downstream effects of SMARCB1 loss depending on differentiation state and identified an interaction between SMARCB1 loss and neural differentiation pressure which causes a resistance to terminal differentiation and a defect in maintenance of a normal cell state. Our results provide insight into how SMARCB1 loss might interact with neural development in the process of ATRT tumorigenesis.

## Introduction

The gene SMARCB1 encodes a subunit of the BAF (also known as SWI/SNF) chromatin remodeling complex. The BAF complex uses ATP hydrolysis to restructure chromatin through alterations of nucleosome positioning and occupancy (Cairns 2007), leading to downstream changes in chromatin accessibility (Tolstorukov et al. 2013; Kadoch et al. 2017) and enhancer activity (Nakayama et al. 2017; Wang et al. 2017). The BAF complex has important roles in development and cellular differentiation. Subunit composition has been shown to change as pluripotent cells differentiate (Lessard et al. 2007; Ho and Crabtree 2010), and a distinct version of the complex with defined subunit composition has been identified in stem cells (Ho et al. 2009). In addition, members of the complex have been identified as reprogramming factors to generate pluripotent cells from somatic cells (Singhal et al. 2010). Nucleosomal occupancy changes are an important aspect of the epigenetic alterations that undergo cellular differentiation (West et al. 2014), and the BAF complex in general along with SMARCB1 in particular have been shown to be important for the regulation of normal nucleosomal occupancy patterns (Tolstorukov et al. 2013; You et al. 2013), with downstream effects on transcription factor binding, enhancer activity and gene expression.

In addition to their normal roles during development, many BAF complex genes have demonstrated roles as tumor suppressor genes. When taken together, the 20 BAF subunit genes have been shown to be mutated in 19% of all tumor types (Shain and Pollack 2013). This speaks to the important genome-wide role of this complex in maintenance of a stable epigenome. Genetic loss-of-function of SMARCB1 in particular has been shown to be both sufficient and necessary for tumorigenesis of atypical teratoid rhabdoid tumors (ATRT) (Versteege et al. 1998; Reincke et al. 2003; Jackson et al. 2009), a highly aggressive and early onset pediatric brain tumor. The mutation rate in ATRT is very low (Lee et al. 2012; Johann et al. 2016), with no other consistent recurrent mutations identified. This low number of mutations is consistent with an early age of onset, but also implies that SMARCB1 loss likely leads to tumorigenesis through initiation of epigenetic changes rather than through the combined effect of multiple genetic mutations. With a median age of onset of 11 months and a lethality rate of 80-90% (Roberts and Orkin 2004), these tumors are responsible for a huge loss of potential life. In addition, very few effective therapies are available for the treatment of ATRT and treatment is complicated by the negative cognitive effects of brain radiation in young children (Ginn and Gajjar 2012). Targeted therapeutics could provide a much-needed alternative to radiation, the development of which would be aided by a greater understanding of the mechanisms driving ATRT tumorigenesis and access to additional model systems with relevance to the human disease.

While transcriptomic and epigenomic analyses of ATRT samples (Johann et al. 2016; Torchia et al. 2016; Chun et al. 2019; Erkek et al. 2019) have characterized the epigenetic alterations which take place following SMARCB1 loss, the mechanisms by which SMARCB1 loss leads to these changes and the factors required for SMARCB1 loss to initiate cellular transformation are not well understood. Increased PRC2 binding (Wilson et al. 2010; Kadoch et al. 2017) and skewed SMARCB1-deficient BAF complex binding at super-enhancers (Nakayama et al. 2017; Wang et al. 2017) have been suggested mechanisms of tumorigenesis due to SMARCB1 loss, but many questions still remain unanswered. The sufficiency of SMARCB1 deletion to drive pediatric tumor growth but lack of SMARCB1 mutation as an exclusive driver mutation in adult cancers, along with the demonstrated role of the BAF complex in development and differentiation leads us to the hypothesis that the ability of SMARCB1 deletion to cause tumorigenesis may be dependent on the epigenetic environment of a particular stage in cellular differentiation. Engineering of induced pluripotent stem cells (iPSCs) with known tumorigenic alterations has been shown to be an effective technique for modeling of glioblastoma (Koga et al. 2020), leading us to apply an inducible system of SMARCB1 loss in iPSCs to address this question.

## Results

### SMARCB1 loss causes differential phenotypes in pluripotent and committed cell types

To interrogate possible interactions between SMARCB1 loss and cellular differentiation state, we generated a doxycycline-inducible SMARCB1 loss of function system in an iPSC line using an inducible shRNA construct targeting SMARCB1 (Fig. 1A, S1A). To rule out the possibility that any observed effects could be due to shRNA off-target effects on genes other than SMARCB1, a doxycycline-inducible SMARCB1 re-expression vector was engineered with either three (m3) or six (m6) silent mutations in the shRNA target sequence (Fig. 1B, S1A). Treatment of this cell line with doxycycline resulted in rapid reduction of SMARCB1 transcript and protein levels (Fig. 1C,D), both of which were successfully rescued in the presence of the re-expression vector. With this inducible system, SMARCB1 loss could be initiated at various stages of differentiation to observe the interplay between cell state and the effects of SMARCB1 loss. After initial doxycycline induction at the iPSC state, it was observed that prolonged induction of SMARCB1 loss resulted in a pronounced cell death phenotype in shSMARCB1 iPSCs (Fig. 1E,F) but not in control iPSCs engineered with a non-targeting shRNA. Beginning three days post-doxycycline induction, a pronounced decrease in growth rate was observed (Fig. 1G) along with an increase in cell death as measured by cell cycle assay, which showed an increase in Sub-G phase dead and dying cells (Fig. 1F). This SMARCB1-induced cell death phenotype is consistent with mouse model data showing embryonic lethality of SMARCB1 knockout mice (Roberts et al. 2000; Han et al. 2016), but has not been previously demonstrated in human cells. Cell death induced by SMARCB1 loss was replicated in a separate doxycycline-inducible SMARCB1 knockdown iPSC cell line utilizing a CRISPR interference method of transcription repression (Fig. S2A-E). However, this system proved to be less stable than the shRNA method and was subject to silencing during differentiation. For this reason, all differentiation experiments were conducted using the shRNA knockdown method with rescue vector. To investigate whether the effects of SMARCB1 loss might vary with differentiation state, iPSCs were differentiated into neural progenitor cells (NPCs) according to the protocol described by Reinhardt et al. (Reinhardt et al. 2013) (Fig. S1B) prior to exposure to doxycycline. Cells were induced with doxycycline for 5 days and monitored for changes in morphology or growth rate. In contrast to the iPSCs, SMARCB1 knockdown NPCs tolerated the loss and displayed no changes in growth rate or morphology (Fig. 1E,H), even with extended doxycycline treatment (data not shown) and a similar level of knockdown as observed in the iPSCs (Fig. S1C). SMARCB1 knockdown NPCs displayed changes in expression of BAF complex subunits similar to those observed in SMARCB1-deficient rhabdoid cell lines and reductions in BAF complex stability (Fig. S1D) consistent with those observed in the literature with ATRT cell lines (Wang et al. 2017), suggesting that shRNA knockdown of SMARCB1 has a similar molecular effect to SMARCB1 loss occurring through genomic deletion.

**Figure 1.**
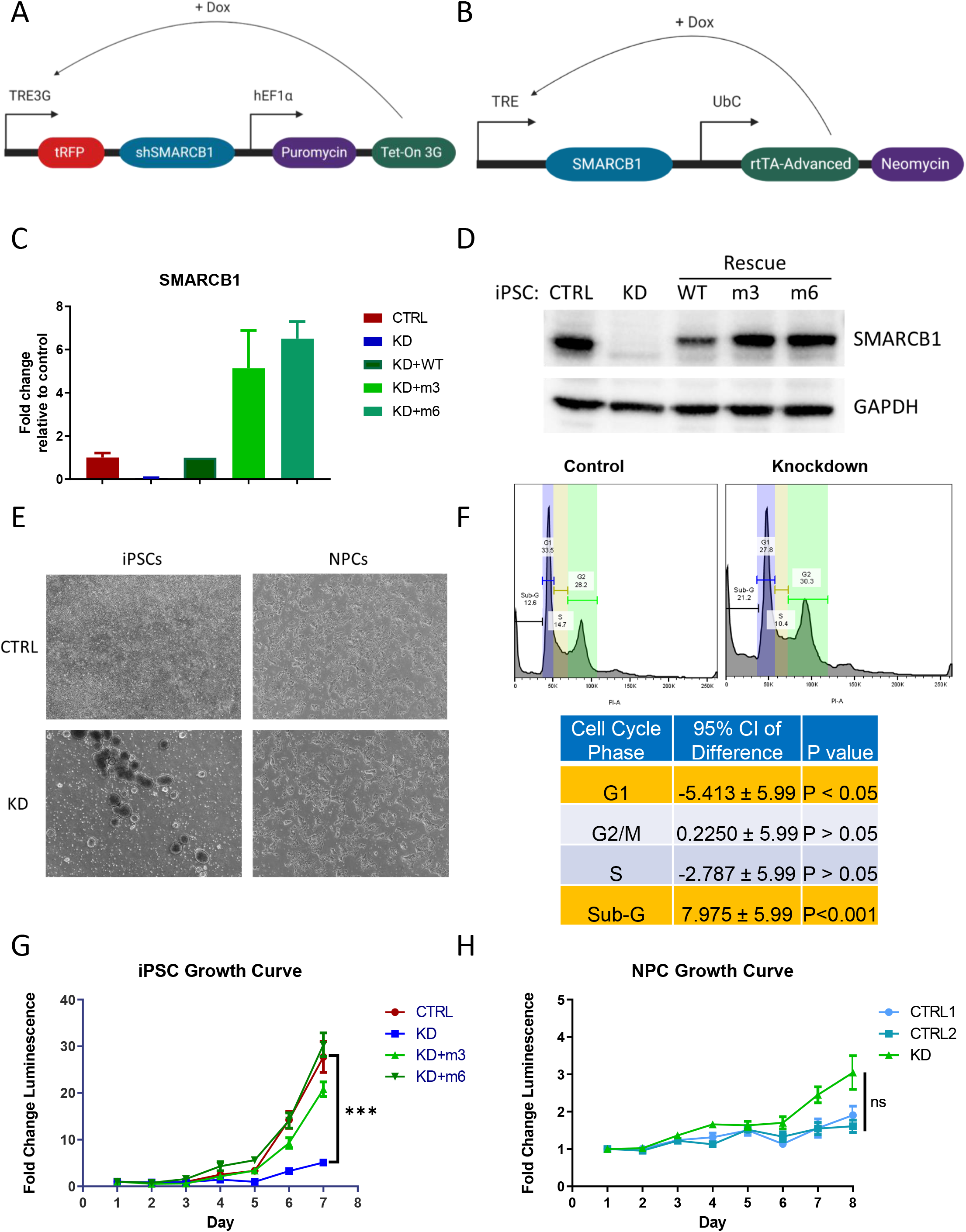
Development of an inducible SMARCB1 knockdown system reveals that SMARCB1 loss causes lethality in pluripotent cells but not neural progenitors. (A) Schematic representation of doxycycline-inducible SMARCB1 shRNA construct, which was stably transduced into induced pluripotent stem cells (iPSCs). (B) Schematic representation of doxycycline-inducible SMARCB1 rescue construct, which was stably transduced into shSMARCB1 iPSCs to rescue SMARCB1 knockdown. Efficacy of shSMARCB1 and rescue vector was tested in iPSCs after 3 days of doxycycline induction using (C) qRT-PCR to measure SMARCB1 transcript levels and standard deviation relative to control mean, or (D) western blot to measure SMARCB1 protein levels. (E) Phase contrast images at 4X magnification of SMARCB1 knockdown iPSCs and NPCs after 5 days of doxycycline induction. (F) Cell cycle assay of 5 day SMARCB1 knockdown iPSCs. Above, FACS readout of PI stained cellular DNA content and corresponding phases of the cell cycle. Numbers indicate percentage of total in each phase. Below, table of percentage differences and 95% confidence interval between control and knockdown at each cell cycle stage. Gold indicates statistically significant differences with 8 replicates. Growth of (G) iPSCs and (H) NPCs was also assessed with doxycycline induction beginning at Day 0 of assay.

To identify transcriptional differences underlying these contrasting phenotypes, we conducted RNA sequencing on control and SMARCB1 knockdown iPSCs and NPCs. In both cell types, more downregulated genes were observed in SMARCB1 knockdown cells than upregulated genes (Fig. 2A, S2D,E). This is consistent with the previously described mechanism for epigenetic and transcriptional changes underlying ATRT, in which loss of SMARCB1 leads to a decrease in BAF complex activity and a corresponding decrease in H3K27Ac active histone marks, along with altered activity of the repressive polycomb repressive complex 2 (PRC2) (Wilson et al. 2010; Kadoch et al. 2017; Nakayama et al. 2017; Erkek et al. 2019). Comparison of the genes differentially expressed by SMARCB1 loss in the two cell types revealed very little overlap between knockdown NPCs and iPSCs (Fig. 2B), suggesting that the downstream targets of SMARCB1 can vary substantially in different cell types. Gene ontology analysis of the dysregulated genes show similarities in the classes of genes altered by SMARCB1 loss in the two cell types, including genes associated with neural development, cellular proliferation and cellular adhesion (Fig. 2C,D). However, many of these shared genes were altered in opposite directions in iPSCs and NPCs, both on the ontology level (Fig. 2E) and on the individual gene level (Fig. 2F). About a quarter of genes which were dysregulated in both iPSCs and NPCs were upregulated in one cell type but downregulated in the other. This unexpected result suggests that the transcriptional effects of SMARCB1 loss can vary dramatically in different epigenetic environments, even leading to opposite phenotypic and transcriptional effects, and explains the very different growth phenotypes observed in knockdown iPSCs and NPCs. These results, along with the established role of the BAF complex in developmental processes (Lessard et al. 2007; Ho et al. 2009; Ho and Crabtree 2010), lead us to believe that SMARCB1 loss might also have dramatic impacts on cellular differentiation processes, potentially highlighting an interplay between differentiation state and ATRT tumorigenesis.

**Figure 2.**
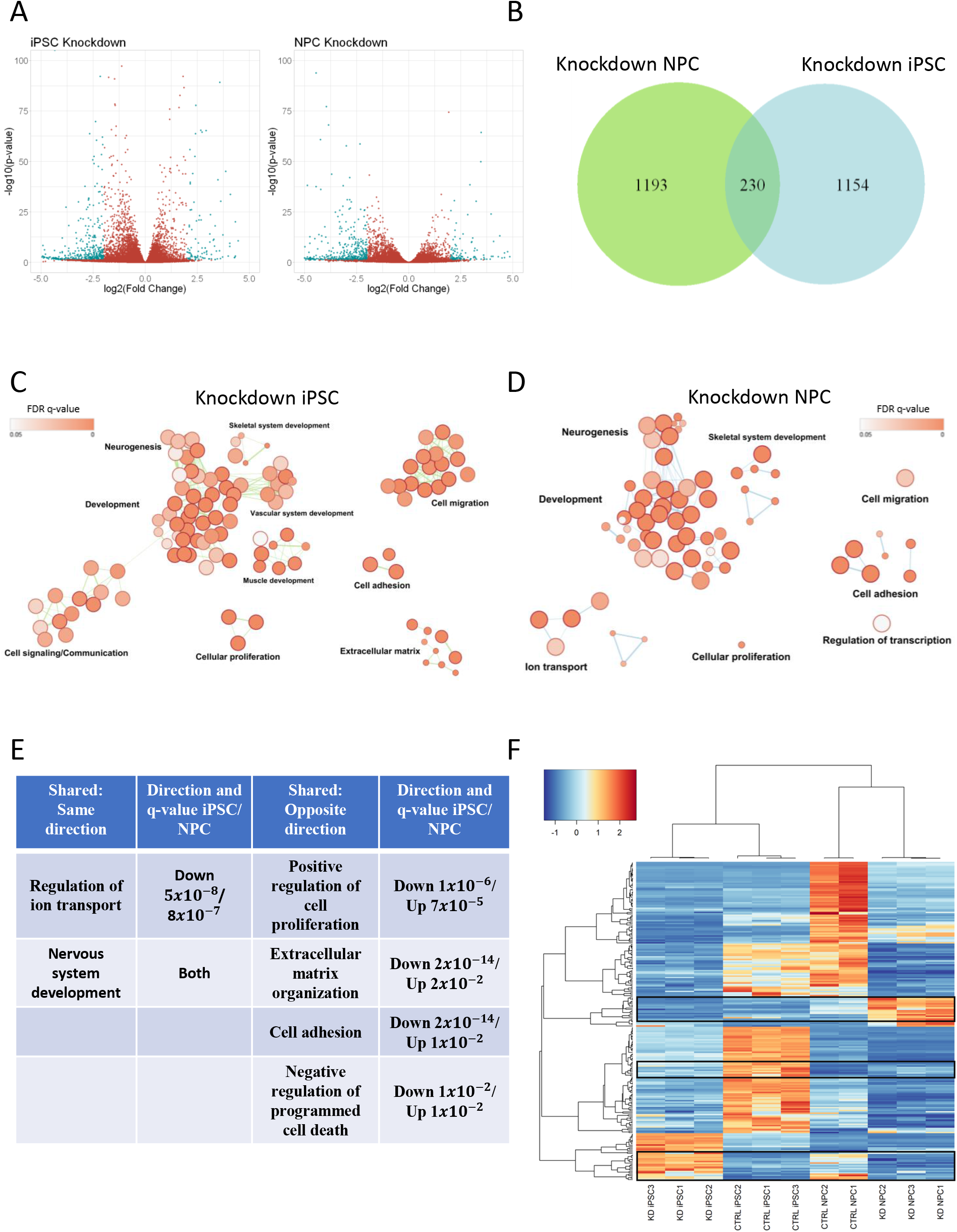
SMARCB1 loss leads to differing transcriptional effects at defined stages of differentiation. RNA sequencing was conducted on SMARCB1 knockdown and shControl iPSCs and NPCs. (A) Volcano plot of genes differentially expressed in SMARCB1 knockdown cells relative to controls in each cell type. (B) Overlap of genes differentially expressed in SMARCB1 knockdown iPSCs and NPCs relative to controls. (C, D) Gene ontology networks for differentially expressed genes in (C) iPSCs and (D) NPCs. Dots represent statistically significant gene ontology terms, clustered based on overlap of the genes contained in each term. Dot size indicates the number of genes included in each term and darker color corresponds to smaller adjusted p-value. Labels indicate the main process making up each cluster. (E) Table comparing shared gene ontology results and direction of alteration between SMARCB1 knockdown iPSCs and NPCs. q-value was obtained using the Benjamini-Hochberg adjustment for multiple comparisons. (F) Heatmap showing genes which are differentially expressed in both iPSCs and NPCs. Boxes indicate regions which are altered in opposite directions between iPSCs and NPCs. Overall, 62 out of 230 overlapping genes (27%) were altered in opposite directions between the two cell types.

### Neural development without SMARCB1 leads to defects in neuron formation in an organoid model

To assess the effect of SMARCB1 loss on neural differentiation, we utilized a cerebral organoid model of neural development (Lancaster and Knoblich 2014) (Fig. 3A). Because this protocol results in the formation of multiple regional identities without selecting for specific neural cell types (Lancaster et al. 2013), the model allows a relatively unbiased assessment of the impact of SMARCB1 loss on the neural developmental process. shControl or shSMARCB1 iPSCs were induced to form cerebral organoids with doxycycline induction beginning at various time points through the protocol and assessed for changes in expression of various neural marker genes (Fig. S3A). While no obvious changes were observed in markers of pluripotency or neural progenitor formation (Nanog, Pax6), decreases were observed in markers of neuronal commitment and maturation (Dcx, visible trend in Map2), especially with earlier doxycycline induction. It was also observed that these early knockdown organoids demonstrated morphological differences relative to the control during expansion and early maturation phases of the protocol (Fig. 3B). Knockdown organoids were defective for the outward expansion of neuroepithelium (Lancaster et al. 2013) into the surrounding matrix typically observed after Matrigel embedding at Day 7 of the differentiation protocol, suggestive of a defect in normal cell differentiation. These results imply that there is a window early in development where cells are especially sensitive to the effects of SMARCB1 loss.

**Figure 3.**
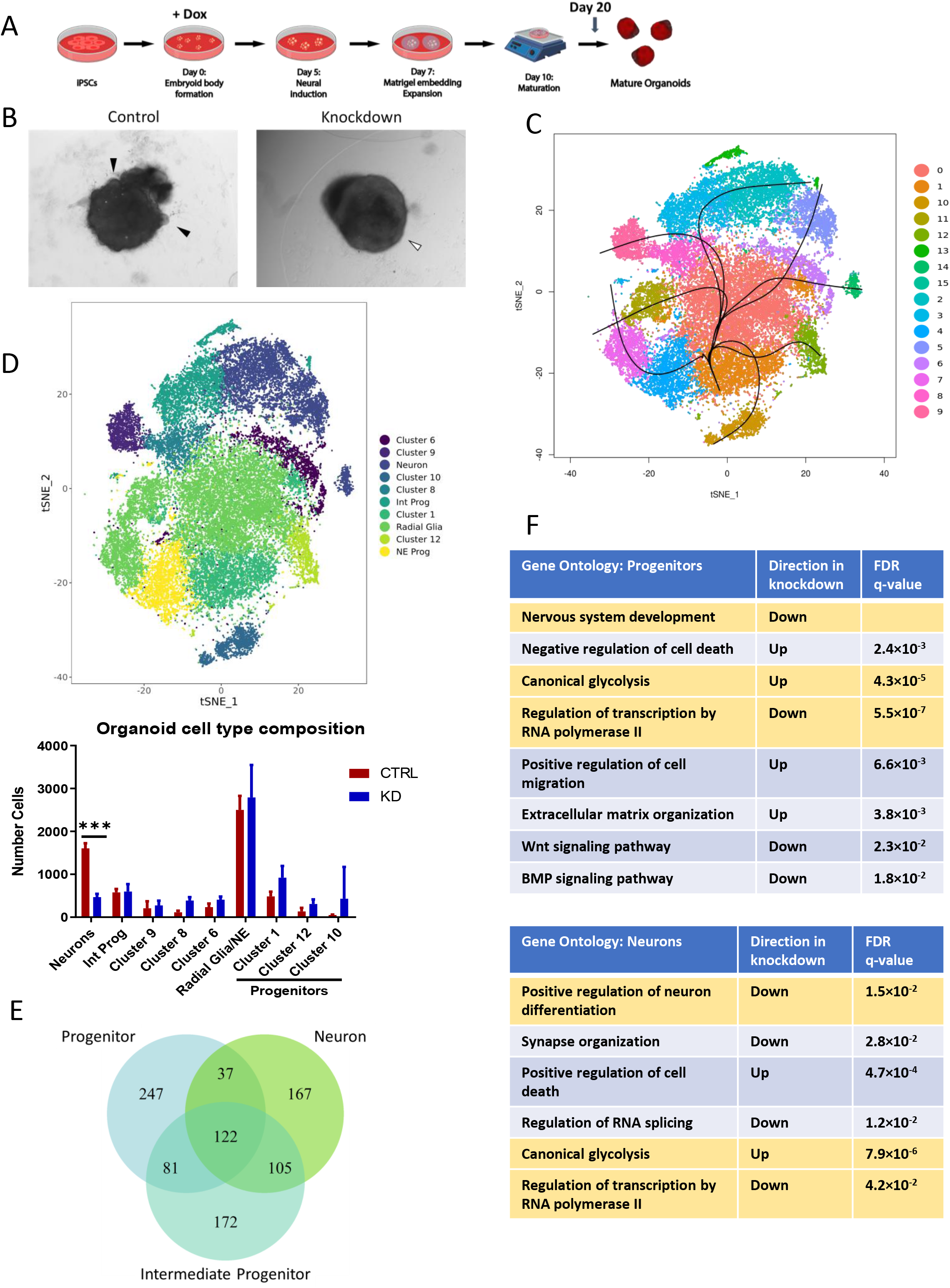
SMARCB1 loss during cerebral organoid development leads to neural differentiation defects. Cerebral organoids were formed from shSMARCB1 and shControl iPSCs in the presence of doxycycline from Day 0 of differentiation protocol. (A) Schematic showing stages of organoid generation from iPSCs. (B) At day 20 of the protocol (10 days of maturation), organoids were examined for morphology and presence of neuroepithelial expansion (black arrows) at 4X magnification. White arrow indicates absence of neuroepithelial expansion. (C) Single-cell RNA sequencing was conducted on three Day 20 control and SMARCB1 knockdown organoids using droplet-based scRNA-seq methodology. Canonical correlation analysis was conducted on combined single-cell data and displayed on a tSNE graph. Clustering and pseudotime analysis was conducted on the combined data to identify variability in cell types and lineages within the organoids. Fifteen clusters were identified, of which cluster 15 was excluded from analysis due to its small size. (D) Left, clusters were analyzed for expression of neural differentiation markers and grouped together by cell type where possible. Clusters not matched to a particular cell type were left unnamed. Right, mean with standard deviation of the number of each cell type in control and knockdown organoids. Statistical comparisons were conducted using two-way ANOVA with Sidak multiple comparisons test. *** indicates adjusted p-value < 0.001. (E) Venn diagram of the overlap across cell types of genes differentially expressed in SMARCB1 knockdown organoids relative to control organoids. (F) Tables showing top gene ontology results from genes differentially expressed in SMARCB1 knockdown organoids relative to controls relative to a list of all expressed genes in progenitor cells, left, and neurons, right. Ontologies highlighted in gold are similarly altered in both cell types.

To better assess the impact of early-stage SMARCB1 loss on neural differentiation we conducted droplet-based single-cell RNA sequencing (scRNA-seq) on three control and three SMARCB1 knockdown organoids at Day 20 of the differentiation protocol, when morphological differences were apparent. These six organoids (Fig. S3B) were aligned and clustered using canonical correlation analysis (Butler et al. 2018) in order to compare numbers of neural cell types between control and knockdown organoids. Cluster analysis resulted in 15 distinct clusters (Fig. 3C, Fig. S3C-D), one of which was excluded for containing less than 100 cells. All but one of these remaining clusters (cluster 10) contained a similar distribution of cells across replicate organoids (Fig. S3B). Organoids had similar distributions of UMI counts and detected genes (Fig. S3E), and SMARCB1 knockdown organoids displayed loss of SMARCB1 transcript in nearly all cells analyzed (Fig. S3F). Clusters were analyzed for expression of several neural development marker genes to identify corresponding cell types (Fig. S4A-F) and Slingshot pseudotime analysis (Street et al. 2018) was performed to identify differentiation trajectories across clusters (Fig. 3C). This analysis revealed a mix of clusters representing neural progenitors, positive for markers such as Sox2, Pax6, and Hes1 (Fig. S4A, C, D), and various stages of neuronal differentiation including intermediate progenitors (Fig. S4E), immature neurons, and more mature neurons (Fig. S4F). Within the progenitor clusters, some seemed to represent neuroepithelial cells and some radial glia (Fig. S4B, D), while others were negative for markers of either of these cell types and may represent progenitors of a distinct lineage (clusters 1, 10, 12). Other clusters were defined by aspects of cell state such as cell cycle stage or apoptosis (Fig. S4B) rather than cell type. Grouping together the identifiable clusters representing neuroepithelial progenitors (cluster 4) and radial glia-like cells (clusters 0, 7, 11), intermediate progenitors (clusters 3, 13), and committed neurons (clusters 2, 5, 14) (Fig. 3D), the number of cells in each group were quantified in both control and SMARCB1 knockdown organoids. The number of neuron-associated clusters was substantially lower in SMARCB1 knockdown organoids (p < 0.001) than controls (Fig. 3D, Fig. S3D) and the expression of individual neuronal markers was lower in knockdown organoids (Fig. S4G), suggesting that the knockdown might be causing a differentiation block and preventing cells from achieving a neuronal cell fate. Although no differences were observed in the number of neuroepithelial or radial glial progenitors, some apparent increases (although not statistically significant to p < 0.05 with current number of replicates), spread across progenitor clusters 1, 12 and 10 (Fig. 3D, S3D) suggest that SMARCB1 loss might lead to a shift in the lineage preference of cells during differentiation while contributing towards a preference for less differentiated cell types.

Differences in the gene expression changes caused by SMARCB1 loss across cell types within the organoids highlight the effects of SMARCB1 loss on differentiation as well as further demonstrating that the transcriptional effects of SMARCB1 loss vary with cell state. For differential expression analysis, related clusters were combined to form larger groups representing different stages of neural differentiation: neural progenitor cells (combining neuroepithelial progenitors, radial glia, and progenitor clusters 1 and 12), intermediate neuronal progenitors, and committed neurons. For each cell type, differential expression analysis was conducted comparing cells of that type in the control and knockdown organoids. A similar number of genes were significantly dysregulated in each cell type, but only about a quarter of these were dysregulated in all three cell types (Fig. 3E). In addition, the number of overlapping genes was greater between more closely related cell types (progenitors and intermediates or intermediates and neurons) than between the more distantly related progenitors and committed neurons. This suggests that there may be a spectrum of transcriptional changes occurring without SMARCB1 that varies throughout the developmental process and has different effects on cells at different stages of cellular differentiation. Gene ontology analysis of dysregulated genes in neural progenitors and neurons showed that different biological processes were affected in the two cell types (Fig. 3F). While canonical glycolysis was upregulated in both cell types and genes associated with both neural development and transcriptional regulation were downregulated, cell death processes were altered in opposite directions. In addition, neural progenitors had additional changes in pathways associated with cellular migration, extracellular matrix organization, Wnt signaling and BMP signaling which were not observed in the more differentiated cells. These differences in transcriptional state illustrate how SMARCB1 loss could lead to distinct cellular phenotypes depending on the cellular epigenetic landscape.

### SMARCB1 loss during neuronal differentiation leads to a lack of stability among neural progenitors which may contribute to tumorigenesis

To validate that the differentiation defects observed in the organoid system are reproducible in other contexts, and to further investigate the effects of SMARCB1 loss on neural differentiation processes, iPSCs were induced with doxycycline and simultaneously differentiated into neural progenitor cells (Fig. 4A). Resulting progenitors were cultured for several passages post-differentiation to assess their ability to maintain an NPC state, and it was observed that NPCs differentiated without SMARCB1 were prone to morphology changes 2-5 passages post-differentiation (Fig. 4A), while control and SMARCB1 rescue NPCs maintained a consistent morphology for up to 10 passages (data not shown). NPCs differentiated without SMARCB1 were also subject to a low-frequency enhancement in growth rate (Fig. S5A), another indication of a lack of stability in these cells relative to control or rescue NPCs. These cells demonstrated a reduction in levels of neural progenitor marker Nestin (Fig. 4B) which is prevented by SMARCB1 rescue, implying a defect in differentiation in the absence of SMARCB1, consistent with observed results using the organoid system. Analysis of BAF complex expression levels in NPCs differentiated without SMARCB1 revealed a decrease in levels of nuclear BAF complex subunits ARID1A, BRG1, SMARCC1, and SMARCD1 relative to control NPCs (Fig. 4C), consistent with what has been observed in SMARCB1 re-expression cell lines (Wang et al. 2017). However, the level of decrease varied substantially in different batches of differentiation for the same level of SMARCB1 knockdown (Fig. 4C), suggesting a stochasticity in the downstream effects of SMARCB1 loss after application of cellular differentiation pressures. RNA-seq of four NPC lines differentiated in the absence of SMARCB1 also revealed a higher transcriptomic variability than was observed in control or rescue cells differentiated with doxycycline or in NPCs subjected to SMARCB1 loss post-differentiation (Fig. 4D). Correlations within replicates of NPCs differentiated without SMARCB1 were significantly lower than rescue NPCs or NPCs with SMARCB1 knockdown induced at the NPC state. A comparison in the genes dysregulated when SMARCB1 is absent throughout NPC differentiation and those altered with SMARCB1 loss at the NPC state (Fig. 4E, S5B,C) revealed that SMARCB1 loss throughout the differentiation process leads to changes in a wide variety of differentiation-associated pathways ranging from renal development to ossification in addition to the expected neural development-associated genes. Changes in pathways associated with cell death, cellular proliferation and TGF-beta signaling are also observed in genes dysregulated by SMARCB1 loss when it occurs during NPC differentiation. A time course of doxycycline application throughout the NPC differentiation process (Fig. S5D) verified that more deleterious effects on neural development are observed with earlier induction of SMARCB1 loss.

**Figure 4.**
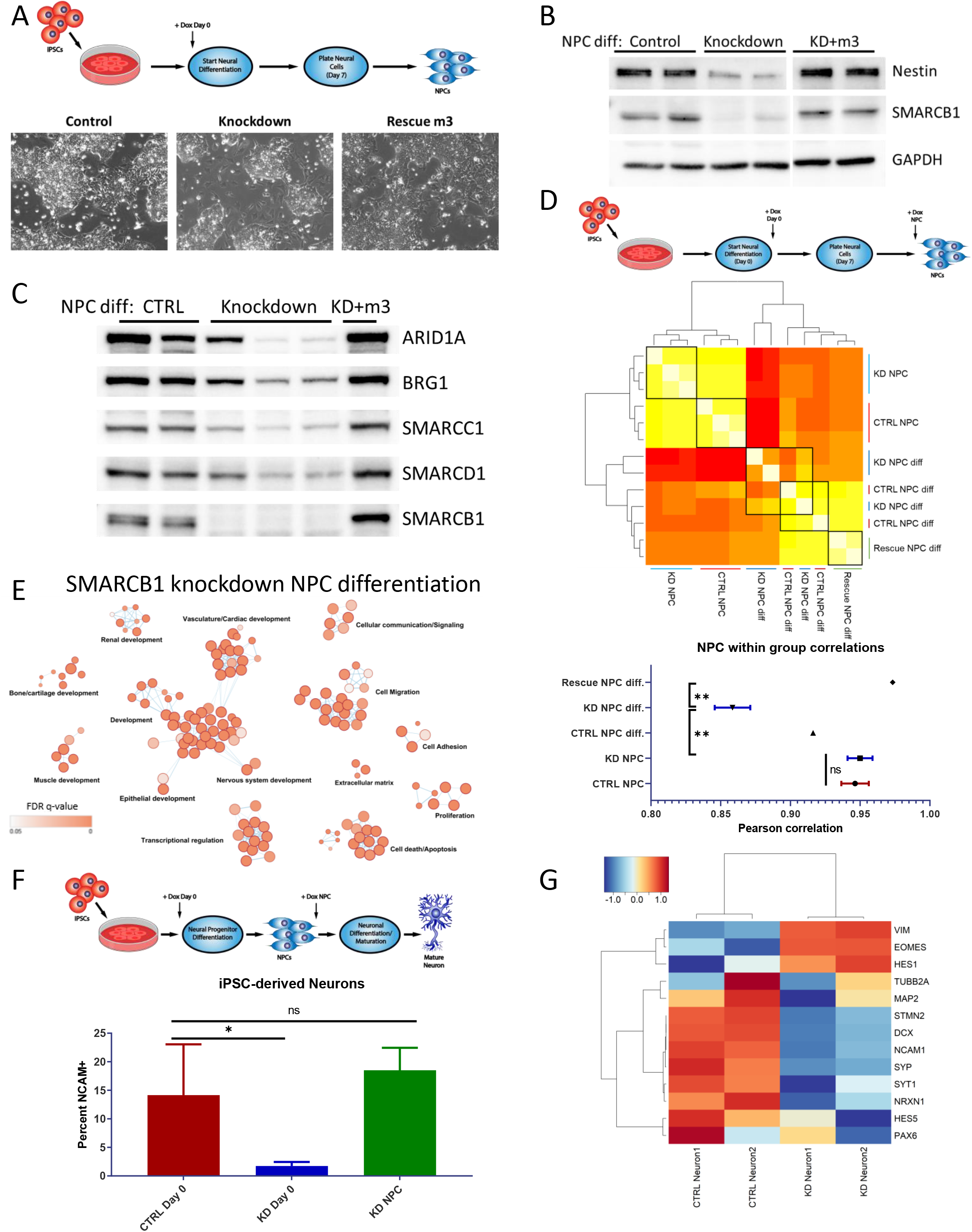
SMARCB1 loss throughout neural differentiation leads to aborted differentiation and a lack of stability in resulting neural progenitor cells. (A) Above, schematic of directed differentiation of iPSCs into neural progenitor cells, with doxycycline induction at Day 0. Below, phase contrast images at 20X magnification of resulting NPC morphology at Day 10-14 of protocol. Cells with abnormal morphology are observed migrating between NPC clusters in cells differentiated without SMARCB1. (B) Western blot showing protein expression of SMARCB1, neural marker Nestin, and control GAPDH in control, SMARCB1 knockdown or rescue NPCs differentiated in the presence of doxycycline. (C) Western blot of BAF complex subunit protein expression in nuclear lysates of NPCs differentiated in the presence of doxycycline. (D) Top, schematic of NPC differentiation with doxycycline induction occurring either at the beginning of the differentiation process (NPC diff.) or at the NPC state (NPC). Middle, Pearson correlation chart comparing transcriptome similarity between control, knockdown or rescue NPCs differentiated with or without doxycycline. Black boxes indicate groups being compared. White corresponds to greater correlation and red to lower correlation. Bottom, mean within group correlation values with standard error of the mean for each group of NPCs. Within group correlations were compared using one-way ANOVA with Tukey’s multiple comparisons test. ** indicates adjusted p-value < 0.01. (E) Gene ontology network of top 500 genes which were differentially expressed in NPCs differentiated with SMARCB1 knockdown relative to controls but not in NPCs with knockdown post-differentiation relative to controls. Dots represent statistically significant gene ontology terms, clustered based on overlap of the genes contained in each term. Dot size indicates the number of genes included in each term and darker color corresponds to smaller adjusted p-value. Labels indicate the main process making up each cluster. (F) Top, schematic showing directed neuronal differentiation with doxycycline induction either at Day 0 or at the NPC state. Control and SMARCB1 knockdown neurons differentiated in this manner were assessed for neuronal maturation efficacy by FACS analysis for neuronal surface marker NCAM1. Bottom, mean and standard deviation percentages of NCAM positive cells in post-differentiation neurons. Comparisons between groups were conducted by one-way ANOVA with Tukey’s multiple comparisons test. * indicates p-value < 0.05. (G) Heatmap of scaled transcript expression of neuronal differentiation markers in control or SMARCB1 knockdown neurons differentiated with Day 0 doxycycline. VIM, HES1, HES5, PAX6, and EOMES are markers of less differentiated neural cells. All other genes are markers of committed neurons.

To validate results from the organoid model and further assess the interaction between SMARCB1 loss and differentiation, control and knockdown cells were subjected to *in vitro* directed neuronal differentiation (Reinhardt et al. 2013). Neuronal maturation efficacy was measured using FACS analysis for surface expression of NCAM, a marker of mature neurons. Cells subjected to SMARCB1 knockdown during both NPC and neuronal differentiation had lower numbers of NCAM positive cells after 25 days of neuronal differentiation and maturation than control cells, as well as when compared to cells subjected to knockdown beginning at the NPC state (Fig. 4F). RNA-seq analysis of control and knockdown neurons showed a reduction in the expression of neuronal markers in cells differentiated in the absence of SMARCB1, along with a retention of some markers of earlier stages of neural differentiation (Fig. 4G) This suggests that SMARCB1 loss during neuronal differentiation leads to a failure in maturation in multiple contexts and validates that cells are particularly vulnerable to SMARCB1 loss early in neural development. This window of vulnerability is consistent between organoid and directed neuronal differentiation experiments, and demonstrates a similar trend to that previously observed in an inducible SMARCB1 knockout mouse model (Han et al. 2016).

### Neural progenitors differentiated without SMARCB1 are transcriptionally similar to ATRT, particularly the SHH subgroup

It seems probable that these observed interactions between SMARCB1 loss and neural differentiation could play a role in ATRT tumorigenesis. To investigate this, previously published bulk RNA-seq data generated from ATRT tumors (Johann et al. 2016) was obtained in order to determine the similarity of this cellular model to patient tumors and to identify cell types with the greatest similarity. To compare the tumor data to the organoid scRNA-seq data, averaged transcriptomic data for each organoid cluster was computed and correlated to the ATRT samples (Fig. 5A). While correlations were generally higher within the organoid or tumor groups, there was variability in the similarity of different organoid cell types to tumors. Neurons in the control organoids were the least similar to the tumors, while progenitor clusters in the SMARCB1 knockdown organoids were most similar (Fig. 5A). This is consistent with the concept of a SMARCB1-deficient early neural progenitor acting as the cell of origin for ATRT. Progenitor clusters in the control organoids were generally less similar to the tumor samples than the same clusters in the knockdown organoids, with the least differentiated clusters (10, 12) showing the greatest similarity to tumors (Fig. 5A). These clusters also demonstrate a possible (but not statistically significant with n=3 organoids) expansion in knockdown organoids relative to controls (Fig. S3D), and thus their development may be favored in the absence of SMARCB1 expression. SMARCB1 knockdown progenitors also show changes in genes associated with transcriptional regulation, nervous system development, and extracellular matrix organization (Fig. 3F), all pathways identified as being altered in ATRT (Johann et al. 2016; Torchia et al. 2016). Comparison of ATRT transcriptomes with RNA-seq data from control and SMARCB1 knockdown iPSCs and NPCs (Fig. 5B-C) revealed greater ATRT similarity to NPCs differentiated without SMARCB1 than either knockdown iPSCs or NPCs induced with SMARCB1 loss post-differentiation. Previous transcriptomic and epigenomic analyses have identified three subgroups within ATRT with differing epigenetic landscapes and gene expression profiles (Johann et al. 2014; Torchia et al. 2016). A comparison of both NPCs differentiated without SMARCB1 via small molecule-directed differentiation and progenitors within SMARCB1 knockdown organoids with ATRTs from each of the three subgroups revealed the greatest similarity with the SHH, or Neurogenic subgroup (Fig. 5D, Fig. S6A-C). This suggests a possible mechanism of ATRT tumorigenesis, likely most relevant to the SHH subgroup, in which focal deletion of SMARCB1 occurs early in neural development, leading to unstable NPCs with tendencies toward differentiation defects, cellular transformation and tumorigenesis (Fig. 6).

**Figure 5.**
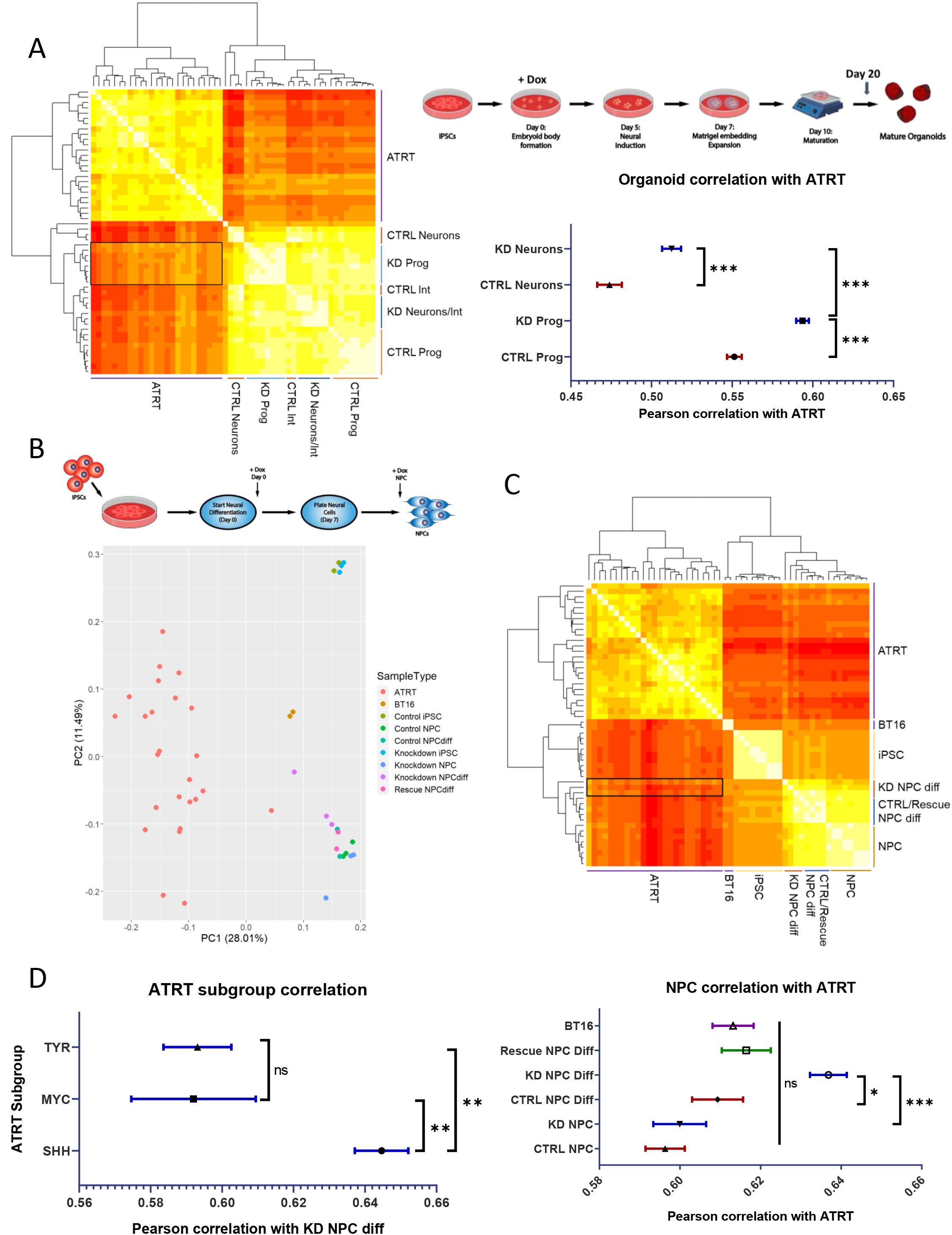
Neural progenitor cells differentiated without SMARCB1 are transcriptomically similar to atypical teratoid rhabdoid tumors. (A) RNA sequencing data from 25 ATRTs (Johann et al. 2014) was compared to averaged single-cell RNA sequencing data for each cluster in control and SMARCB1 knockdown cerebral organoids. Left, chart of pearson correlation values between individual clusters and ATRT samples, clustered by similarity. White indicates highest correlation and red corresponds to lowest correlation. Labels indicate cell types corresponding to clusters. Box indicates region of highest similarity to ATRT samples. Upper right, schematic of protocol used for organoid generation. Lower right, mean and standard error of pearson correlation values of progenitors and neurons from control and SMARCB1 knockdown organoids. Comparisons between groups conducted using one-way ANOVA with Tukey’s multiple comparisons test. *** indicates adjusted p-value < 0.001. (B) Top, schematic of protocol used for NPC differentiation either in the presence of doxycycline from Day 0 (NPCdiff) or at the NPC state (NPC). Bottom, principal component analysis of RNA sequencing results from 25 ATRT samples compared to directed differentiation of control or SMARCB1 knockdown iPSC-derived NPCs, along with BT16 ATRT cell line and undifferentiated iPSCs induced with doxycycline. (C) Top, chart of pearson correlation values between control and SMARCB1 knockdown iPSCs and NPCs, induced with doxycycline during and post-differentiation, along with ATRT samples, clustered by similarity. White indicates highest correlation and red corresponds to lowest correlation. Box indicates region of highest similarity to ATRT samples. Bottom, mean and standard error of pearson correlation values of control and SMARCB1 knockdown NPCs and BT16 cell line with ATRT samples. Comparisons between groups conducted using one-way ANOVA with Tukey’s multiple comparisons test. * indicates adjusted p-value < 0.05 and *** indicates adjusted p-value < 0.001. (D) Mean and standard error of pearson correlation values of NPCs differentiated without SMARCB1 and samples from each of the three ATRT subgroups. Comparisons between groups conducted using one-way ANOVA with Tukey’s multiple comparisons test. ** indicates adjusted p-value < 0.01.

**Figure 6.**
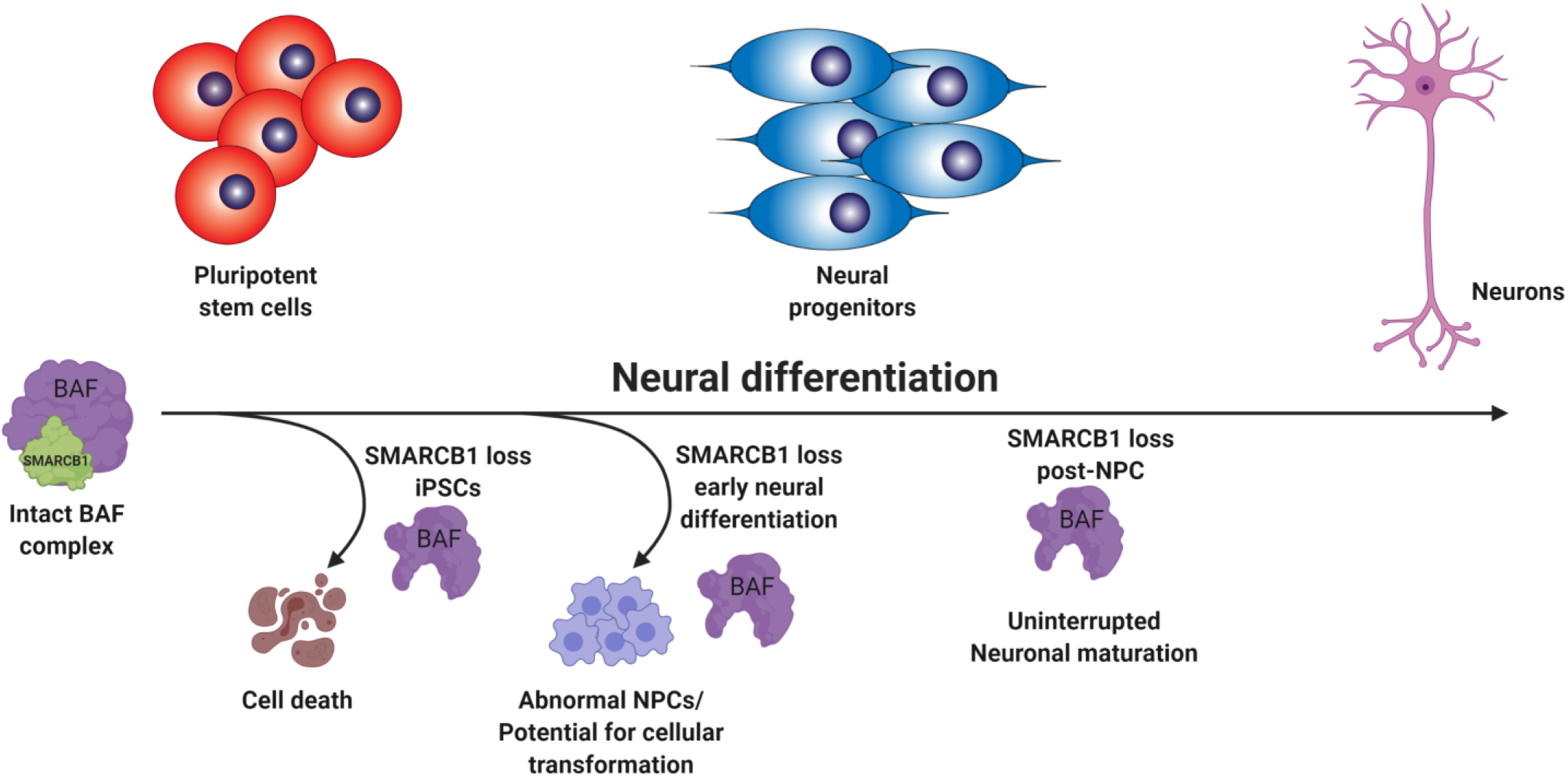
SMARCB1 loss interacts with developmental state to redirect cell fate. Schematic summarizing findings on the interaction between neural differentiation state and the effect of SMARCB1 loss. In pluripotent cells, SMARCB1 loss results in cell death. In the early stages of neural differentiation, SMARCB1 loss induces dedifferentiation, morphology changes and lack of stability in resulting NPCs along with defects in capacity for further neuronal differentiation. With induction of knockdown in later stages of differentiation, little to no effect on differentiation capacity or cell growth was observed.

## Discussion

SMARCB1 is an important chromatin remodeling subunit as well as a known tumor suppressor whose loss is the primary driver of pediatric rhabdoid tumors. In this study we have interrogated the interactions between SMARCB1 loss, cellular differentiation state, and transcriptional changes associated with tumorigenesis, while generating a cellular model which will have utility for future mechanistic studies as well as for identification of potential therapeutic vulnerabilities in SMARCB1-deficient cells. While other systems of SMARCB1 loss or reintroduction have been used to study the mechanisms underlying ATRT in a controlled manner (Wilson et al. 2010; Han et al. 2016; Nakayama et al. 2017; Wang et al. 2017; Carugo et al. 2019; Langer et al. 2019), this complementary system has the benefit of using human cells, having the flexibility to take into account the effects of differentiation processes, and using SMARCB1 loss alone without additional oncogenic drivers, consistent with the human tumor phenotype (Lee et al. 2012). In addition, similar to a recent publication (Langer et al. 2019), our study provides an interrogation of the interactions between SMARCB1 loss and neural development, however, here we illustrate novel insight into the dramatic phenotypic differences which can occur with loss of SMARCB1 at different stages of differentiation, such as lethality in pluripotent cells and impairment of neuronal commitment and maturation. This is the first study to model the interaction between SMARCB1 loss and cellular differentiation state that likely contributes to ATRT tumorigenesis in human cells, and to monitor the accompanying gene expression and phenotypic changes.

We have demonstrated significant differences in the response of cells to SMARCB1 loss at differing stages of neural differentiation and identified a window early in neural commitment in which cells seem to be particularly vulnerable to SMARCB1 loss of function and in which SMARCB1 loss results in profound defects in the progression of differentiation. SMARCB1 loss during this period results in cells with greater similarity to ATRT tumors than loss at an earlier pluripotent or later committed neural progenitor state, along with a lack of stability resulting in a tendency toward stochastic alterations in cellular morphology and gene expression. This provides insight into a possible mechanism for ATRT tumorigenesis in which a loss of SMARCB1 during embryogenesis could result in cells which are primed for cellular transformation. This is consistent with both the early age of onset of this disease and heterogeneity of presentation, as well as with mouse data showing development of ATRT-like tumors with SMARCB1 loss during early mouse embryogenesis (Han et al. 2016). While it is clear that the neural progenitors differentiated without SMARCB1 in this study are most similar to the SHH/Neurogenic subgroup of ATRT, more work is needed to determine the mechanism underlying this similarity. It is possible that the transcriptomic and epigenetic differences between the subgroups are driven by different cells or developmental stages of origin, and the origin of SHH/Neurogenic tumors more closely resembles the loss of function early in development which was applied in this study. Indeed, the SHH/Neurogenic subgroup of tumors does tend to occur in younger children, consistent with this hypothesis (Johann et al. 2016; Torchia et al. 2016). Another possibility is that the mechanism of SMARCB1 loss could play a role, with the larger chromosomal alterations observed more often in the other subgroups of ATRT leading to additional effects on neighboring genes or regulatory regions not replicated with a knockdown model. Thus the predominance of smaller focal or point mutations in the SHH subgroup might more closely resemble a SMARCB1 knockdown system.

In this study we focused on the interactions of SMARCB1 loss with neural development, but molecular heterogeneity and dysregulated developmental pathways observed in extra-cranial malignant rhabdoid tumors (Chun et al. 2016; Chun et al. 2019) suggest that a similar mechanism might take place in other types of rhabdoid tumors. In all, we have presented an in depth investigation into the stages of neural differentiation in which SMARCB1 loss has the greatest effects on cellular outcome, chronicled gene expression changes resulting from SMARCB1 loss at various stages of differentiation and generated a novel platform on which to expand our understanding on the mechanisms and vulnerabilities underlying ATRT tumorigenesis.

## Methods

### Pluripotent stem cell culture and neural differentiations

Human induced pluripotent stem cell (iPSC) line iPS12 was purchased from Cell Applications. This cell line is integration-free and was validated for pluripotency, viability, karyotype normality and normal disease status by Cell Applications. CRISPRi Gen1C and WTC iPS lines were obtained from the Conklin lab at University of California San Francisco. iPSCs were cultured using standard feeder-free conditions with mTESR1 or mTESR Plus medium on Matrigel-coated plates. iPSCs were induced to form neural progenitor cells using a small-molecule based differentiation protocol as described in Reinhardt et al. (Reinhardt et al. 2013), using combined small-molecule inhibition of BMP and TGFβ signaling along with WNT and SHH pathway stimulation. Neuron differentiations were also conducted as described in Reinhardt et al. for peripheral neurons (Reinhardt et al. 2013), starting from NPCs of 3-6 passages in smNPC maintenance medium (N2B27 medium + CHIR + PMA). Neurons were harvested after two weeks in neuronal maturation medium (N2B27 medium + dbcAMP + TGF-b3 + BDNF + GDNF). Both neural progenitor and neuron differentiations were conducted under 0.5 μg/mL puromycin to prevent loss of shRNA expression. Rescue cell lines were differentiated in the presence of 0.5 μg/mL puromycin and 100 μg/mL G418. When needed, doxycycline was applied at a 1 μg/mL concentration for all experiments.

### Plasmids and cell line engineering

Doxycycline-inducible shRNA constructs against SMARCB1 and non-targeting controls were purchased from Dharmacon (SMARTvector Inducible Lentiviral shRNA) and transductions were conducted using Dharmacon Trans-Lentiviral Packaging Kit according to kit protocol. After selection with puromycin, individual clones were screened for SMARCB1 knockdown by quantitative real-time PCR and top clones were pooled to obtain highly efficient knockdown. Of three shRNA constructs tested, only one was capable of efficient SMARCB1 knockdown (sh905). Rescue vectors were engineered from pInducer20 plasmid backbone (Addgene #44012) to express SMARCB1 cDNA with either 3 or 6 silent mutations at the sh905 target site. See supplementary figure 1 for shRNA target and rescue sequences. For CRISPRi experiments, CRISPRi Gen1C cell line was transduced with several guide RNAs targeting the SMARCB1 locus and selected with blasticidin for guide RNA expression. Clones were then obtained, screened for SMARCB1 knockdown and top clones pooled to obtain a highly efficient level of knockdown in iPSCs.

### Organoid development and culture

Organoids were generated using the STEMdiff Cerebral Organoid Kit from Stemcell Technologies (Cat. 08570), which is based on the Lancaster et al.(Lancaster and Knoblich 2014) protocol for cerebral organoid formation and development. Organoids were developed in the presence of 0.5 μg/mL puromycin and 1 μg/mL doxycycline. Organoids were matured for 10-50 days in Maturation Medium (Day 10 of protocol). Organoids used for single-cell RNA-seq were matured for 10 days to look at early-stage commitment and development of neural progenitors.

### Growth, cell cycle and cell death assays

For growth assays, 1000-2000 cells/well with 5-10 replicates per cell line were plated on Matrigel-coated black 96-well plates in maintenance medium without antibiotic selection. First timepoint was read within 24 hours of plating for baseline comparison and subsequent readings were performed every 24 hours following. Medium changes were conducted as needed (every 2-3 days) throughout the assay. ATPlite 1step assay kit (PerkinElmer 6016731) was used to estimate cell number. For cell cycle assays, eight replicate doxycycline inductions of ~1×10^6^ cells were harvested and fixed overnight in 70% ethanol, washed three times with PBS and stained for 30 minutes with 0.5 mL FxCycle PI/RNase Staining Solution (Life Technologies F10797) before quantification with a BD LSR II flow cytometer. Cell cycle percentages were calculated using FlowJo software and the Dean-Jett-Fox model. Doxycycline was added to shRNA iPSCs for 5 days before fixation and to CRISPRi iPSCs for 9 days before fixation.

### Western blots and immunoprecipitations

For western blots, cells were lysed in RIPA lysis buffer with proteinase and phosphatase inhibitors. 20 μg lysate was run on an SDS-PAGE gel, transferred at 350 mA over 1.5 hours to EMD Millipore Immobilon-P PVDF membrane, blocked for one hour in 5% BSA, and probed with primary antibody overnight. Membranes were washed with PBS + 0.1% Tween-20, probed for 1-2 hours with secondary HRP-conjugated antibody and exposed using Thermo Scientific SuperSignal West Pico or Femto Chemiluminescent Substrate. For BAF complex immunoprecipitations, nuclear extractions were first performed using Thermo Scientific NE-PER Nuclear and Cytoplasmic Extraction Reagents (78833) according to kit instructions. For immunoprecipitations, 2 μg of BRG1 antibody (Santa Cruz, sc-17796) was incubated for one hour with 20 μL Dynabeads Protein G magnetic beads (Thermo Fisher Scientific, 10004D), washed, then incubated overnight at 4 °C with 500 μg of nuclear lysate, washed and prepared for SDS-PAGE. Washes were conducted with either citric or RIPA buffer, as specified. Western blots were run as previously described. Primary antibodies used for western blots were: SMARCB1/BAF47 (mouse, 1:500, BD Biosciences, 612110), GAPDH (rabbit, 1:5000, 2118), HDAC1 (rabbit, 1:1000, Cell Signaling, 2062), SMARCD1 (mouse, 1:1000, Santa Cruz, sc-135843), SMARCC1 (rabbit, 1:1000, Santa Cruz, sc-10756), BRG1 (mouse, 1:1000, Santa Cruz, sc-17796), Nestin (rabbit, 1:1000, EMD Millipore).

### Bulk RNA sequencing preparation and analysis

RNA was extracted using Qiagen RNeasy Plus Mini Kit and library prep was conducted using Illumina NEBNext Ultra Directional Library Prep Kit. Samples were sequenced using an Illumina sequencer with a minimum of 20 million reads per sample. Transcriptome data was aligned using the STAR aligner to a reference human genome (hg19). Reads were counted using featureCounts with default settings, and differential expression analysis conducted using DESeq2 R package (Love et al. 2014). Significant genes were considered to be those with a Benjamini-Hochberg adjusted p-value of greater than 0.05 and a fold-change of greater than 2. Gene ontology analysis for upregulated and downregulated genes was conducted using the GOrilla web-based tool (Eden et al. 2009), comparing a list of up to 500 most significant genes (based on adjusted p-value) to a background list of all expressed genes (rpkm > 4 across all samples). For gene ontology networks, top 500 significant genes were analyzed for GO biological process and Reactome biological pathway enrichment by gProfiler (Raudvere et al. 2019) and output file visualized using Cytoscape (Shannon et al. 2003) software with EnrichmentMap plugin (Merico et al. 2010).

### Single-cell RNA sequencing preparation and analysis

Three replicate control and knockdown Day 20 organoids (10 days maturation) were collected, washed and incubated for one hour on a 37 °C shaker in Accutase + 50 μg/mL DNaseI to aid in generation of a single-cell suspension. Organoids were dissociated by gentle pipetting with a wide-bore pipette after 15 and 45 minutes of incubation. Clumps were removed by filtration through a cell strainer, resuspended in PBS+0.04% BSA, counted and resuspended to a 1.2×10^6^ cells/mL concentration. Cell viabilities were 60-80% with a total of 5×10^4^ to 2×10^5^ cells collected per organoid. Samples were prepared for single-cell RNA sequencing as detailed in the 10X Chromium Single Cell 3’ Reagent Kits v2 User Guide. Reagents used included the Chromium Single Cell 3’ Library & Gel Bead Kit v2 (PN-120237), Chromium Single Cell A Chip Kit (PN-120236) and Chromium i7 Multiplex Kit (PN-120262). The protocol was followed for a desired 10,000 cells per organoid using 10 cycles of cDNA amplification. Quality control was conducted after cDNA amplification and library construction on a BioAnalyzer TapeStation instrument. Sample libraries were pooled, and shallow sequencing conducted on an Illumina HiSeq4000 to estimate cell numbers and read counts for each sample and a new pool generated to obtain 50000 reads/cell for each sample. Final sequencing was conducted on an Illumina NovaSeq 6000. Analysis of the resulting data was conducted using the Cell Ranger pipeline for counting and aggregation of sequencing reads. Additional analysis was then conducted using the Seurat R toolkit for single cell genomics (Butler et al. 2018). Cells with a mitochondrial DNA percentage above 0.1 were filtered out of subsequent analysis. Canonical correlation analysis was conducted using variable genes from the control and knockdown conditions, and a tSNE plot of the first 20 aligned subspaces used for visualization. Clusters were generated using a resolution value of 0.6 and the first 20 CCA subspaces, resulting in the identification of 15 clusters, of which cluster 15 was excluded from later analysis due to its small size. Identification of cluster markers, differential expression analysis, and cluster quantifications were all conducted using the Seurat toolkit. Gene ontology analysis of differential expression gene sets was conducted using the Gorilla (Eden et al. 2009) website in comparison to a background list of expressed genes.

### Flow cytometry

For neuron FACS experiments, Day 25 neurons were dissociated with Accutase + DNase I for 30 minutes, with occasional gentle pipetting to break up clumps. Cells were filtered through a cell strainer and resuspended in PBS + 1% FBS. 1×10^6^ cells were stained with NCAM-1 antibody (CD56 Anti human Alexafluor 700, 1:200 dilution, Fisher Scientific, BDB557919) for 1 hour, washed three times with PBS + 1% FBS, and analyzed on a Sony SH800 instrument.

## Acknowledgements

We would like to thank Dr. Bruce Conklin for providing CRISPR interference cell lines and constructs and Dr. Diana Hargreaves for provision of BAF complex antibodies. In addition, we would like to thank Rachel Reed and Drs. Ciro Zanca, Jorge Benitez, Jianhui Ma, Amy Haseley Thorne, Nathan Jameson and Tiffany Taylor for their technical contributions to this project and Drs. Gene Yeo, Robert Wechsler-Reya, Rafael Bejar, Alysson Muotri, and Lorraine Pillus for their continued advice and mentorship.

This work was supported by Padres Pedal the Cause/RADY grant (#PTC2017), the National Institute of General Medical Sciences T32GM008666 (A.P.), the National Institute of Neurological Diseases and Stroke R01-NS080939 (F.F.), and a Japan Society for the Promotion of Science (JSPS) Overseas Research Fellowship (S.M.). This publication includes data generated at the UC San Diego IGM Genomics Center utilizing an Illumina NovaSeq 6000 that was purchased with funding from a National Institutes of Health SIG grant (#S10 OD026929).

## Author contributions

AP performed cell line generation, all analytical experiments and sample processing, genomic analysis and played a primary role in drafting of the manuscript. TK aided in cell line generation and maintenance and single-cell RNA sequencing sample preparation in addition to aiding in initial project conception and data interpretation. SM aided in cell culture experiments and provided guidance to the project. PDJ and MK collected and analyzed ATRT RNA sequencing data and provided them for use in this project. JRC aided in obtaining funding for this project and provided advice. FF provided funding for the project, was responsible for the conception of the work and contributed to data interpretation and project design. All authors contributed to revising of the manuscript.

