## Supplemental Figures for "SMARCB1 loss interacts with neuronal differentiation state to block maturation and impact cell stability"

A

| Construct | Sequence | RNA region targeted | SMARCB1 region |
|---|---|---|---|
| shSMARCB1 | TTCAAATCCAGATCGTCAC | ORF | Exon 6 |
| shSMARCB1 target site (WT rescue) | GTGACGATCTGGATTTGAA | | |
| shSMARCB1 m3 rescue | GCGATGACCTCGACTTAAA | | |
| shSMARCB1 m6 rescue | GCGATGACCTCGACTTAAA | | |

B

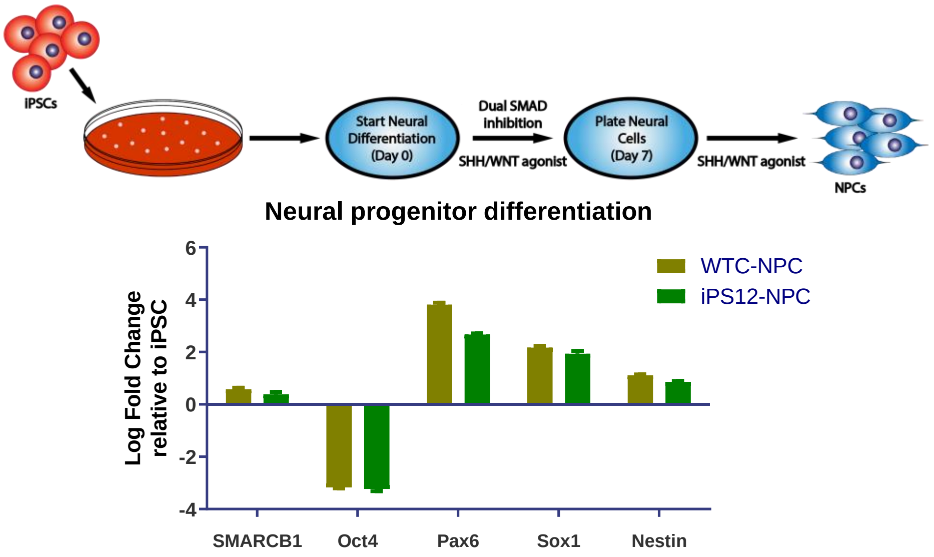


C

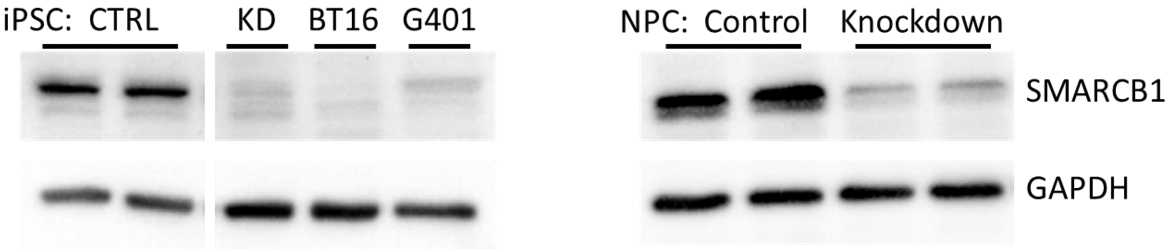


D

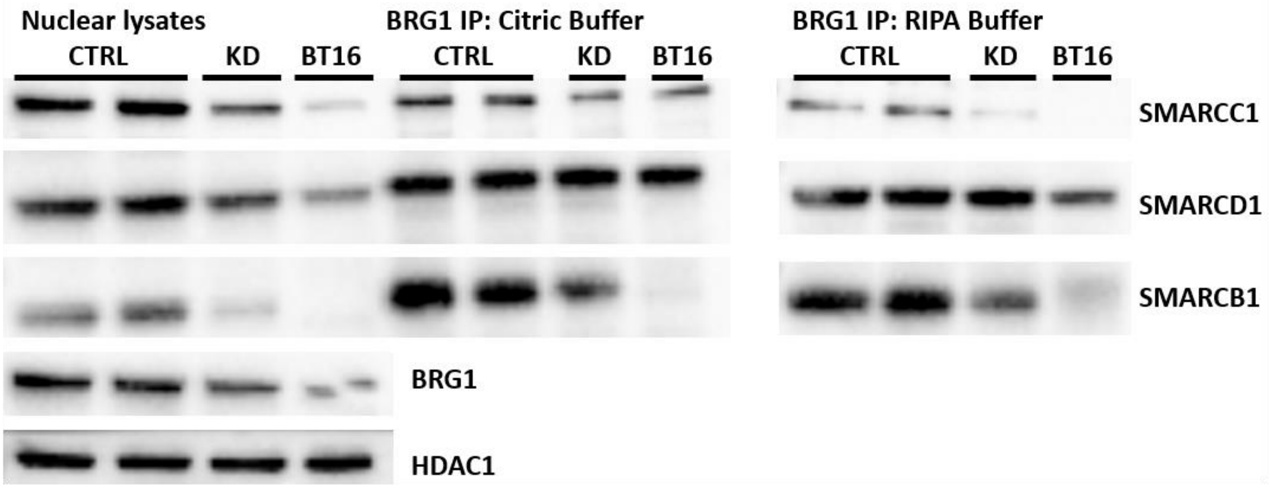

**Supplemental Information**

**Figure S1. Doxycycline-inducible SMARCB1 knockdown system efficiently reduces SMARCB1 levels in both iPSCs and NPCs.** (A) Table of shRNA sequence and target sequence on SMARCB1, along with sequences of rescue vectors at shSMARCB1 target site. Induced mutations for resistance to shSMARCB1 are shown in red. (B) Above, schematic of NPC differentiation protocol and below, qRT-PCR mean and standard deviation of shControl NPC transcript levels of several markers of pluripotency or NPC differentiation relative to those in undifferentiated iPSCs for two different iPSC cell lines. (C) Western blots of SMARCB1 and GAPDH protein levels in iPSCs (left) in comparison to ATRT cell line BT16 and rhabdoid cell line G401 and (right) in NPCs induced for three days with doxycycline. (D) Immunoblot of BAF complex subunits in shControl and SMARCB1 knockdown NPC and BT16 cell line nuclear lysates and BRG1 immunoprecipitation. BRG1 immunoprecipitations were conducted with both a milder citric buffer wash and the more stringent RIPA buffer to assess differences in complex stability under both conditions. HDAC1 in nuclear lysates serves as loading control.

A

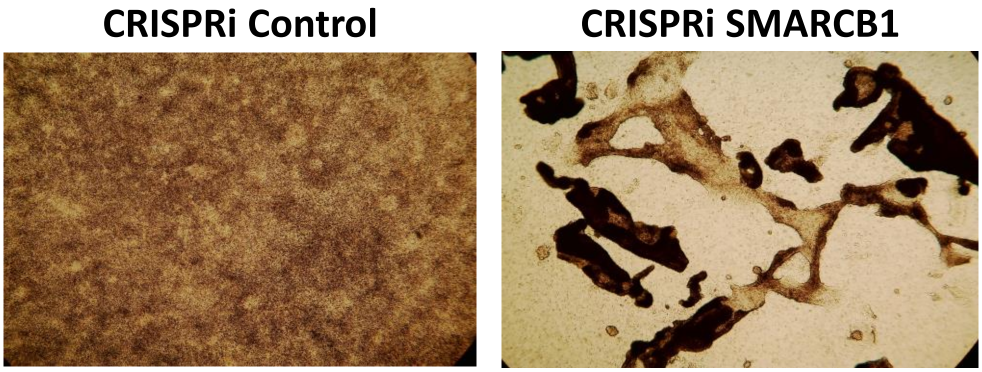


B

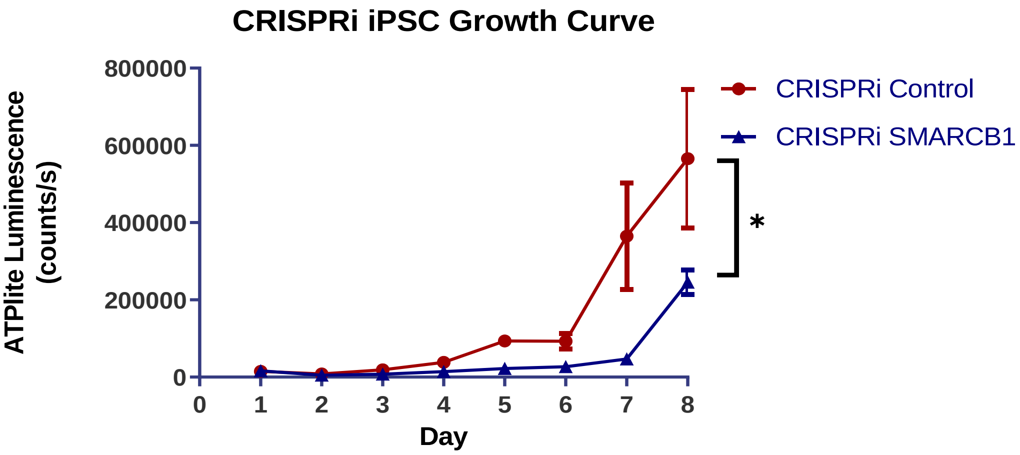


C

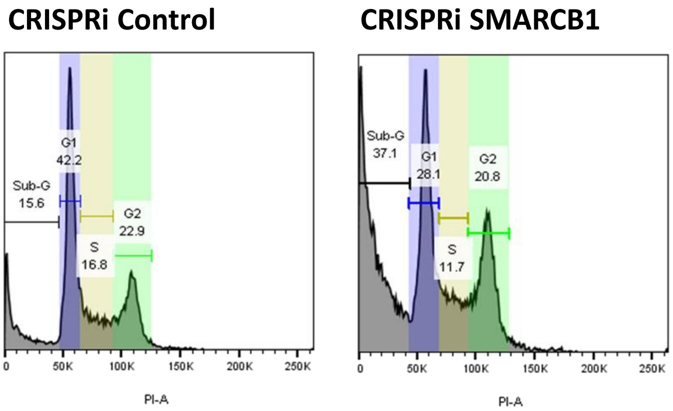


| Cell Cycle Phase | 95% CI of Difference | P value |
|---|---|---|
| G1 | -13.90 ± 9.68 | P<0.001 |
| G2/M | -9.233 ± 9.68 | P < 0.05 |
| S | -0.5467 ± 9.68 | P > 0.05 |
| Sub-G | 23.57 ± 9.68 | P<0.001 |

D

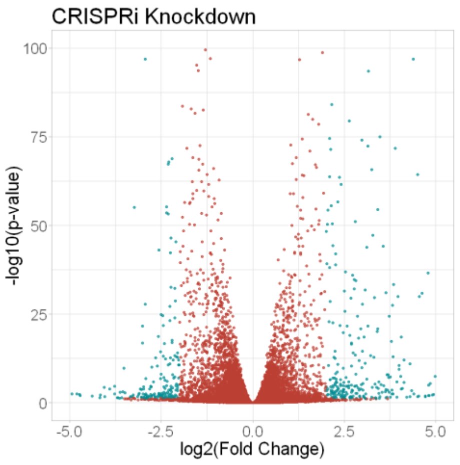


E

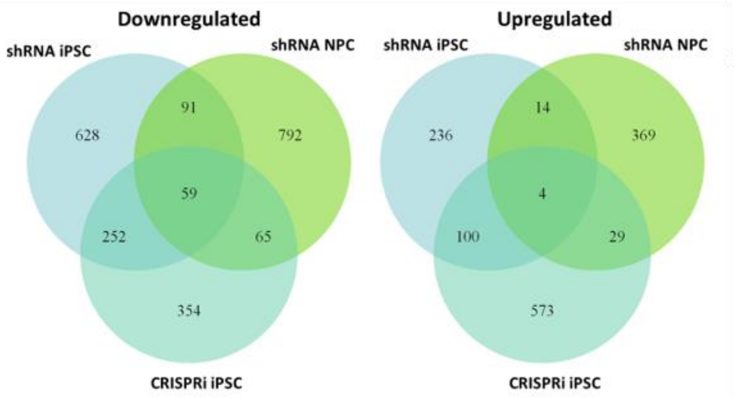

**Figure S2. SMARCB1 knockdown using CRISPR interference shows similar phenotypes and growth effects to shRNA knockdown in iPSCs.** (A) Brightfield images at 2X magnification of cell morphology in CRISPRi control and SMARCB1 knockdown iPSCs after 8 days of doxycycline induction. (B) ATPlite growth curve showing CRISPRi control and SMARCB1 knockdown iPSCs in the presence of doxycycline. * indicates p-value of final timepoint <0.05. (C) Cell cycle assay of CRISPRi control and SMARCB1 knockdown iPSCs. Left, FACS readout of PI stained cellular DNA content and corresponding phases of the cell cycle. Numbers indicate percentage of total in each stage. Right, table of percentage differences and 95% confidence interval between control and knockdown at each cell cycle phase. Gold indicates statistically significant differences with 8 replicates. (D) Volcano plot of SMARCB1 knockdown CRISPRi iPSC RNA sequencing data relative to control. Statistically significant differences are highlighted in teal. (E) Diagram showing overlap of genes differentially expressed in SMARCB1 knockdown cells relative to control in shRNA iPSCs, CRISPRi iPSCs and shRNA NPCs. Greatest overlap is observed between shRNA and CRISPRi iPSCs.

Parisian_Supp_Fig_3

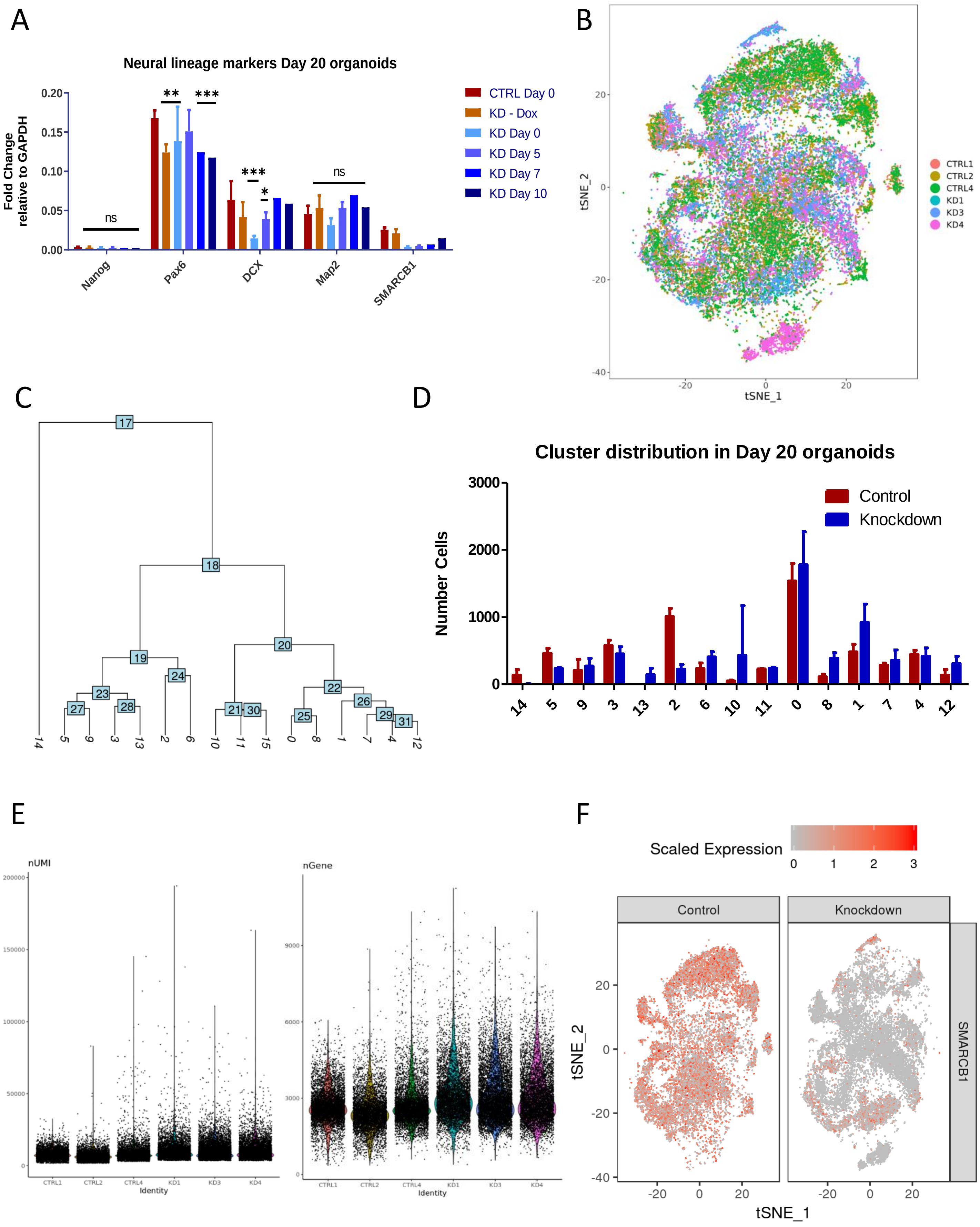

**Figure S3. Additional visualizations and metrics of organoid single-cell RNA-seq data.** (A) qRT-PCR analysis of SMARCB1 transcript levels and neural development-associated genes in Day 20 organoids with doxycycline induction at various timepoints during the organoid development protocol. Plotted is transcript fold-change relative to GAPDH and standard deviation. Comparisons were conducted using one-way ANOVA with Tukey's multiple comparisons test. *** indicates adjusted p-value < 0.001, ** indicates adjusted p-value <0.01, * indicates adjusted p-value < 0.05. (B) Overlaid single-cell RNA sequencing data of three control and SMARCB1 knockdown organoids graphed on a tSNE plot, colored by organoid of origin. (C) Phylogenetic tree of organoid clusters. Clusters on the left represent more differentiated neurons or neuronal intermediates, while those on the right represent less differentiated progenitors. (D) Quantification of the mean and standard deviation number of cells in each cluster in control and SMARCB1 knockdown organoids. (E) Graphs of single-cell RNA sequencing UMI counts and gene counts for individual organoids. (F) tSNE plots of control organoids (left) and SMARCB1 knockdown organoids (right) colored by level of SMARCB1 expression.

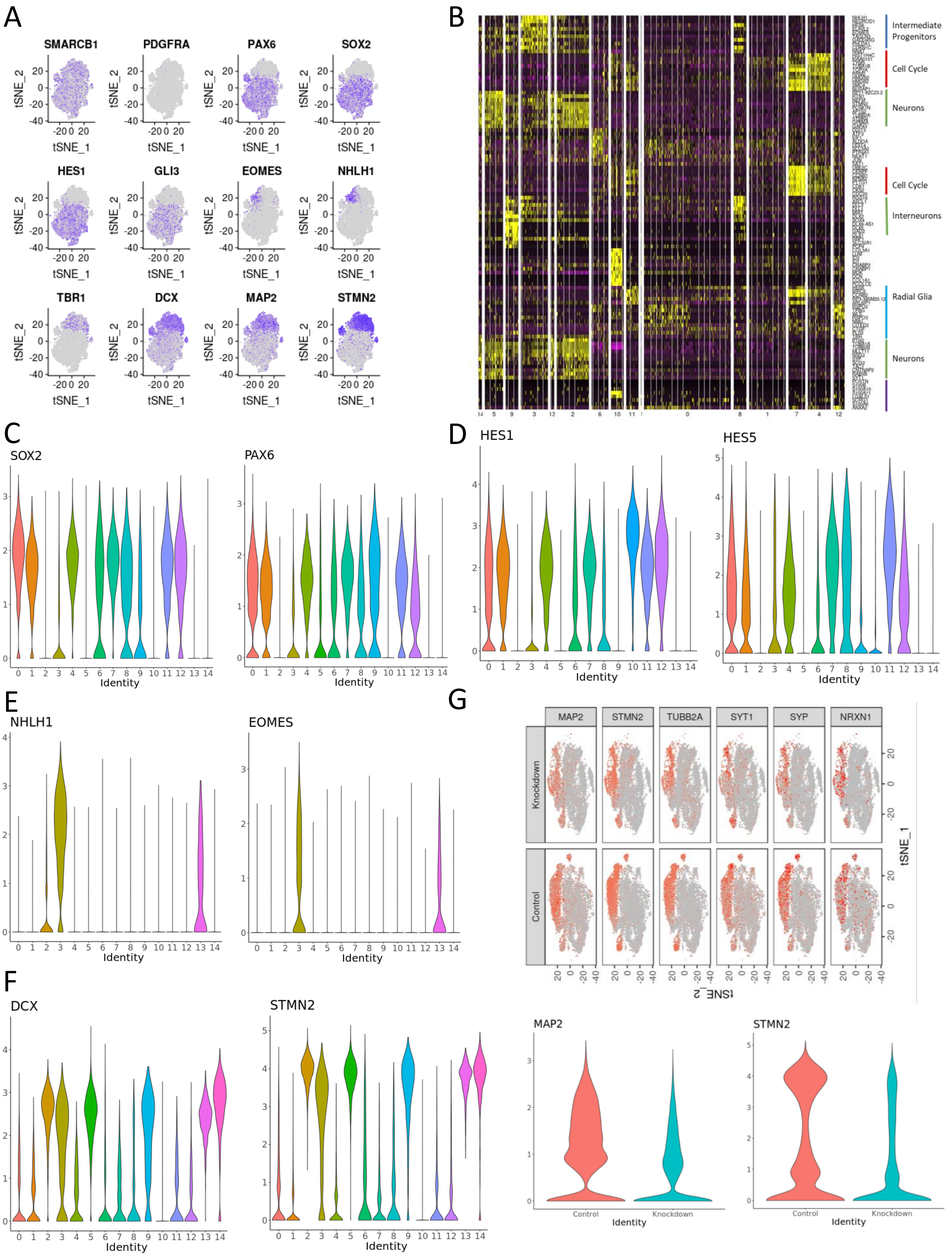
A
SMARCB1
PDGFRA
PAX6
SOX2
HES1
GLI3
EOMES
NHLH1
TBR1
DCX
MAP2
STMN2
tSNE_1
tSNE_2
B
Intermediate Progenitors
Cell Cycle
Neurons
Cell Cycle
Interneurons
Radial Glia
Neurons
C
SOX2
PAX6
Identity
D
HES1
HES5
Identity
E
NHLH1
EOMES
Identity
F
DCX
STMN2
Identity
G
MAP2
STMN2
TUBB2A
SYT1
SYP
NRXN1
Knockdown
Control
tSNE_1
tSNE_2
MAP2
STMN2
Control
Knockdown
Identity

**Figure S4. Cell type determination of clusters in organoid single-cell RNA-seq data.** (A) tSNE plots of overlaid control and SMARCB1 knockdown organoids colored for expression of SMARCB1 and various neural marker genes ranging from those common in less differentiated cells (top) to those common in more differentiated neurons (bottom). (B) Heatmap of top 10 differentially expressed genes in each cluster after a differential expression analysis comparing all clusters. Yellow indicates higher expression and purple/black indicates lower expression. Labels on right indicate cell types or processes enriched in that group of genes. (C) Violin plots showing expression of neural progenitor markers Sox2 and Pax6 across organoid clusters. (D) Violin plots showing expression of intermediate neuronal progenitor markers EOMES and NHLH1 across organoid clusters. (E) Violin plots showing expression of neuronal markers DCX and STMN2 across organoid clusters. (F) Violin plots showing expression of radial glia markers Hes1 and Hes5 across organoid clusters. (G) Above, tSNE plots of control (lower subplot) and knockdown (upper subplot) organoids colored for level of expression of neuron markers MAP2, STMN2, TUBB2A, SYT1, SYP, and NRXN1. Below, violin plots of neuron markers MAP2 and STMN2 in control and knockdown organoids. Knockdown organoids show a lower amount of expression for neuronal markers than control organoids.

A

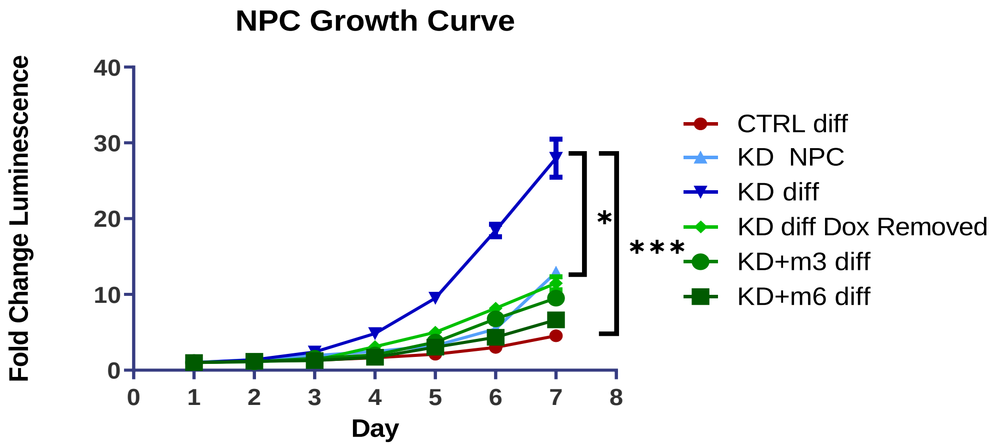
NPC Growth Curve
Fold Change Luminescence
40
30
20
10
0
0 1 2 3 4 5 6 7 8
Day
*
***
CTRL diff
KD NPC
KD diff
KD diff Dox Removed
KD+m3 diff
KD+m6 diff

B

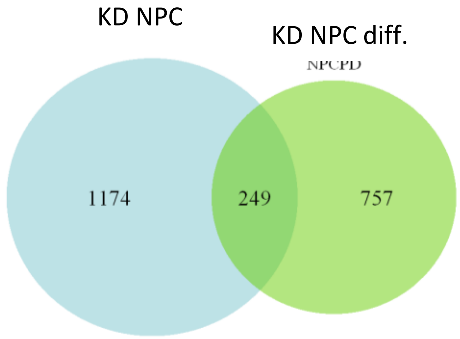
KD NPC
KD NPC diff.
NPCPD
1174
249
757

C

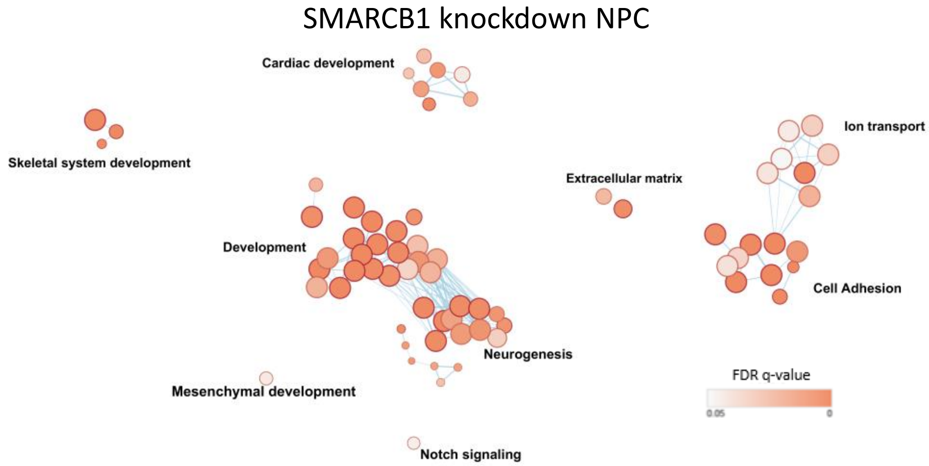
SMARCB1 knockdown NPC
Cardiac development
Skeletal system development
Ion transport
Extracellular matrix
Development
Cell Adhesion
Neurogenesis
Mesenchymal development
FDR q-value
0.05
0
Notch signaling

D

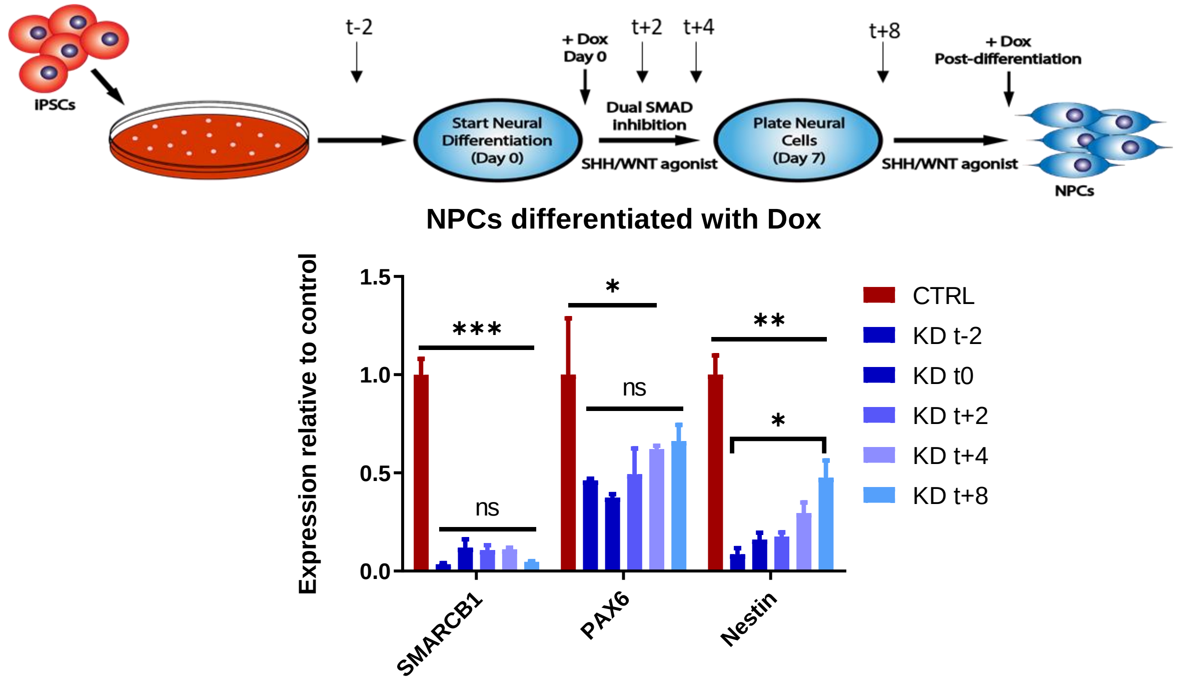
IPSCs
t-2
+ Dox
Day 0
t+2
t+4
t+8
+ Dox
Post-differentiation
Start Neural
Differentiation
(Day 0)
Dual SMAD
Inhibition
SHH/WNT agonist
Plate Neural
Cells
(Day 7)
SHH/WNT agonist
NPCs
NPCs differentiated with Dox
Expression relative to control
1.5
1.0
0.5
0.0
***
ns
*
ns
**
*
SMARCB1
PAX6
Nestin
CTRL
KD t-2
KD t0
KD t+2
KD t+4
KD t+8

**Figure S5. NPCs differentiated without SMARCB1 are prone to changes in morphology and may demonstrate enhanced proliferation or dependency on continued SMARCB1 loss**. (A) ATPlite growth curve of one differentiation batch of control, rescue and SMARCB1 knockdown NPCs differentiated with doxycycline (Day 0) compared to induction with doxycycline post-differentiation (NPC). * indicates a p-value < 0.05, *** indicates a p-value < 0.001. This phenotype was not observed in all batches of differentiation but illustrates a flexibility in NPCs differentiated without SMARCB1 to display unusual changes in morphology or phenotype. (B) Diagram comparing genes differentially expressed in SMARCB1 knockdown condition relative to control in NPCs differentiated without SMARCB1 (NPC diff.) and NPCs induced with doxycycline post-differentiation (NPC). (C) Gene ontology network of genes unique to SMARCB1 knockdown at the NPC state and not altered in NPCs differentiated without SMARCB1. Dots represent statistically significant gene ontology terms, clustered based on overlap of the genes contained in each term. Dot size indicates the number of genes included in each term and darker color corresponds to smaller adjusted p-value. Labels indicate the main process making up each cluster. (D) Above, schematic showing time-course experiment with doxycycline induction at various time points throughout the NPC differentiation process. Below, qRT-PCR analysis showing transcript levels and standard deviation relative to control mean for SMARCB1 and neural progenitor markers Pax6 and Nestin. Upper bars indicate most conservative significance levels between control and knockdown. Lower bars indicate significance between knockdown timepoints. Comparisons were conducted using two-way ANOVA with Tukey's multiple comparisons test. * indicates adjusted p-value < 0.05, ** indicates adjusted p-value < 0.01, *** indicates adjusted p-value < 0.001.

A

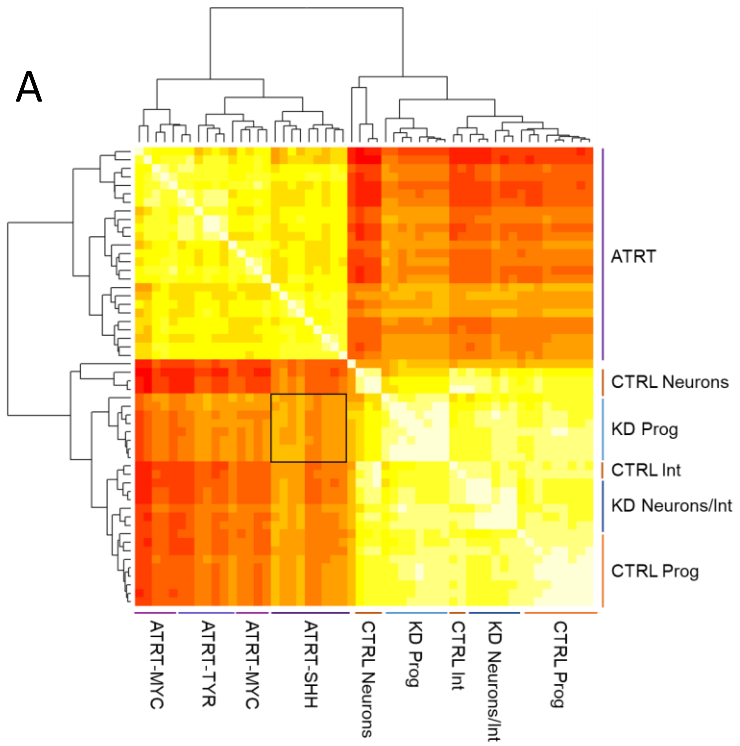
ATRT
CTRL Neurons
KD Prog
CTRL Int
KD Neurons/Int
CTRL Prog
ATRT-MYC
ATRT-TYR
ATRT-MYC
ATRT-SHH
CTRL Neurons
KD Prog
CTRL Int
KD Neurons/Int
CTRL Prog

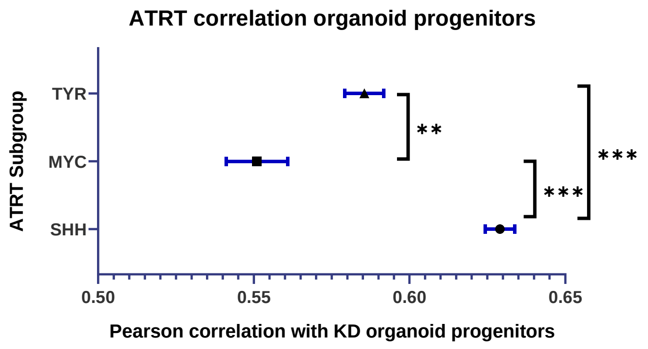
ATRT correlation organoid progenitors
ATRT Subgroup
TYR
MYC
SHH
**
***
***
0.50
0.55
0.60
0.65
Pearson correlation with KD organoid progenitors

B

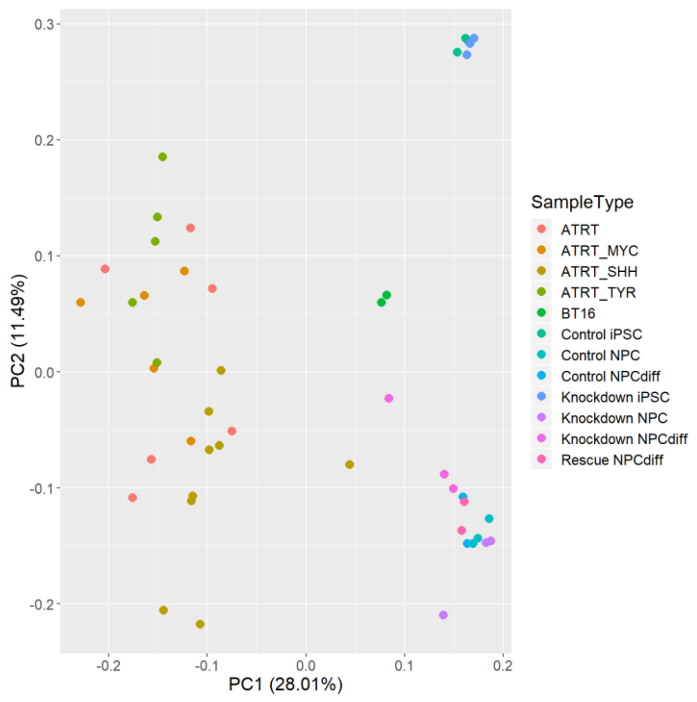
0.3
0.2
0.1
0.0
-0.1
-0.2
PC2 (11.49%)
-0.2
-0.1
0.0
0.1
0.2
PC1 (28.01%)
SampleType
ATRT
ATRT_MYC
ATRT_SHH
ATRT_TYR
BT16
Control iPSC
Control NPC
Control NPCdiff
Knockdown iPSC
Knockdown NPC
Knockdown NPCdiff
Rescue NPCdiff

C

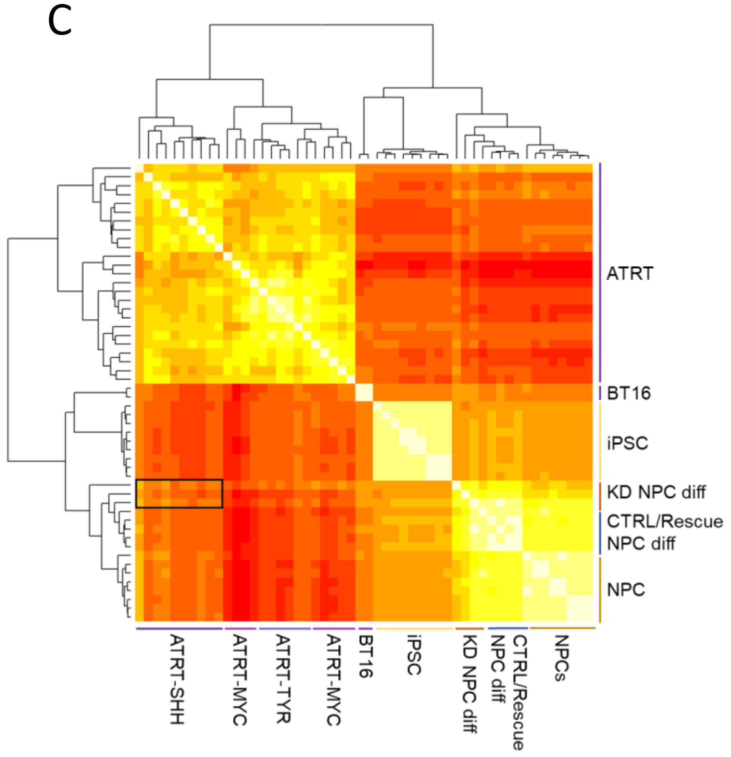
ATRT
BT16
iPSC
KD NPC diff
CTRL/Rescue NPC diff
NPC
ATRT-SHH
ATRT-MYC
ATRT-TYR
ATRT-MYC
BT16
iPSC
KD NPC diff
NPC diff
CTRL/Rescue
NPCs

**Figure S6. NPCs differentiated with SMARCB1 knockdown are most similar to the SHH/Group 1 subtype of ATRT in both organoid and directed differentiation models.** (A) RNA sequencing data from 25 ATRTs was compared to averaged single-cell RNA sequencing data for each cluster in control and SMARCB1 knockdown cerebral organoids. Left, chart of pearson correlation values between individual clusters and ATRT samples, clustered by similarity and labeled by subgroup designation (Johann et al. 2014). White indicates highest correlation and red corresponds to lowest correlation. Labels indicate cell types corresponding to clusters. Box indicates region of highest similarity between ATRT subgroup and organoid clusters. Right, mean and standard error of pearson correlation values of progenitors from SMARCB1 knockdown organoids with ATRTs from each subgroup. Comparisons between groups conducted using one-way ANOVA with Tukey's multiple comparisons test. *** indicates adjusted p-value < 0.001. ** indicates p-value < 0.01. (B) Principal component analysis of RNA sequencing results from 25 ATRT samples, labeled by corresponding subgroup where known, compared to directed differentiation of control or SMARCB1 knockdown iPSCs differentiated into NPCs in the presence of doxycycline (NPCdiff) or at the NPC state (NPC), along with BT16 ATRT cell line and undifferentiated iPSCs induced with doxycycline. (C) Chart of pearson correlation values between control and SMARCB1 knockdown iPSCs and NPCs, induced with doxycycline during and post-differentiation, along with ATRT samples, clustered by similarity and labeled by subgroup designation (Johann et al. 2014). White indicates highest correlation and red corresponds to lowest correlation. Box indicates region of highest similarity between ATRT subgroup and SMARCB1 knockdown model.
